# An Integrated Transcriptomic Landscape of Lung Cancer Identifies Tumor Clusters of Biological Significance

**DOI:** 10.64898/2026.03.16.712177

**Authors:** Sonali Arora, Laavan Suresh, H. Nayanga Thirimanne, Gregory Glatzer, Matt Jensen, Jackson P. Fatherree, Eric Q. Konnick, Kevin M. Levine, Angela N. Brooks, A. McGarry Hougton, Colin C. Pritchard, David MacPherson, Alice H. Berger, Eric C. Holland

**Author notes:** Corresponding author Eric C Holland Human Biology, Fred Hutch Cancer Center, 1100 Fairview Ave N, Seattle, WA 98109.

## Abstract

Lung cancer encompasses multiple histological entities with substantial molecular heterogeneity that remain incompletely resolved at population scale. Here, we constructed a unified reference landscape of lung cancer by analyzing raw RNA sequencing data from 1,824 tumors spanning adenocarcinoma (n=966), squamous cell carcinoma (n=628), small cell lung cancer (n=150), and unclassified non–small cell lung cancer (n=80). Following batch correction, samples were analyzed using consensus clustering and visualized with PaCMAP to generate a molecular atlas annotated with clinical and biological metadata. Rather than segregating by pathological diagnosis, tumors organized along conserved transcriptional axes defined by tumor-intrinsic biology including proliferative or metabolic programs and immune-infiltrated states. Consensus clustering resolved nine robust molecular clusters, including an adenocarcinoma-associated subgroup, a neuroendocrine-like adenocarcinoma marked by ASCL1 activation, immune-associated regions, and bifurcation of both small cell and squamous carcinomas into biologically distinct states. Spatially restricted expression of selected clinically relevant transcripts nominated state-specific therapeutic hypotheses requiring future functional and clinical validation. Projection of patient tumors and patient-derived xenografts onto the atlas demonstrated preservation of transcriptional identity and enabled quantitative assessment of model fidelity. This integrated framework organizes lung cancer as a structured continuum of transcriptional states and provides a reference resource for biological interpretation and future translational studies.

## INTRODUCTION

Lung cancer is broadly classified into two major types: non-small cell lung cancer (NSCLC) and small cell lung cancer (SCLC). NSCLC accounts for approximately 85% of all lung cancer cases and is further subdivided into several histological subtypes, the most common being lung adenocarcinoma (LUAD) and lung squamous cell carcinoma (LUSC). Among these, lung adenocarcinoma is the most prevalent histological subtype of lung cancer and exhibits profound molecular and clinical heterogeneity. It arises from glandular epithelial cells and encompasses diverse oncogenic drivers, transcriptional programs, and tumor microenvironmental states. While subsets such as EGFR-or KRAS-driven adenocarcinomas have been extensively characterized, a substantial fraction of tumors defy clear molecular stratification, complicating prognosis and therapeutic decision-making^1,2^. This heterogeneity suggests that adenocarcinoma is not a single disease entity but rather a collection of biologically distinct states occupying a continuous molecular spectrum^3^.

Small cell lung cancer (SCLC) and non–small cell lung cancer (NSCLC) are traditionally distinguished by histopathology and clinical behavior. A subset of SCLC is characterized by rapid proliferation, neuroendocrine features, and near-universal inactivation of TP53 and RB1, whereas NSCLC represents a heterogeneous category encompassing multiple epithelial lineages and oncogenic mechanisms. Emerging evidence indicates that these categories are not strictly separable at the molecular level, with transcriptional overlap and phenotypic plasticity observed across diagnostic boundaries, particularly between SCLC and adenocarcinoma^4,5^.

Squamous cell carcinoma (SCC) constitutes a distinct major subtype of lung cancer, typically associated with significant smoking exposure and defined by alterations in TP53, SOX2 amplification, and dysregulation of squamous differentiation programs. Despite its relative molecular coherence compared to adenocarcinoma and SCLC, SCC itself exhibits internal heterogeneity, including immune-related and metabolic subtypes, suggesting additional unresolved structure within this histological class^1,6^.

Unified molecular landscapes derived from bulk RNA sequencing provide an unbiased framework for resolving such heterogeneity. By integrating large numbers of previously published datasets, these approaches enable cost-effective reuse of existing data, increase statistical power through substantially larger sample sizes, and capture continuous biological variation that is often obscured by discrete classification schemes. While single-cell technologies offer unparalleled cellular resolution, their limited cohort sizes and experimental cost constrain their ability to map population-level structure, whereas large-scale bulk transcriptomic integration excels at identifying robust, reproducible disease states^7,8^.

The approach presented here builds directly on a series of reference molecular landscapes previously developed by our group for brain tumors^9^, including comprehensive atlases of medulloblastoma, ependymoma^10^, and meningioma^11^. These studies demonstrated that large-scale, batch-corrected integration of bulk transcriptomic data can resolve clinically meaningful disease subgroups, reveal lineage relationships obscured by histopathology, and provide durable community resources for tumor classification and biological discovery. Importantly, these landscapes have been widely adopted as reference frameworks, enabling external investigators to contextualize new samples, validate biomarkers, and generate testable hypotheses without the need for additional sequencing. Extending this paradigm to lung cancer, with its pronounced histological diversity and clinical burden, represents a natural and necessary progression toward a unified, population-scale view of thoracic tumor biology.

In this study, we assembled a large-scale transcriptomic cohort comprising 1,824 lung cancer samples, including adenocarcinoma (n=966), squamous cell carcinoma (n=628), small cell lung cancer (n=150), and unclassified non–small cell lung cancer (n=80), by collecting and uniformly processing raw sequencing data from multiple public sources. Following stringent quality control and batch correction, we constructed an integrated reference landscape using consensus clustering, and visualized using Pairwise Controlled Manifold Approximation (PaCMAP)^12^, enabling joint visualization and analysis of tumors across histological boundaries. We systematically overlaid clinical, demographic, and molecular annotations and applied consensus clustering to define robust disease regions with shared biological programs. This integrated landscape, available at Oncoscape (Lung Landscapes), provides an interactive framework for biomarker discovery, lineage-state interpretation, immune state annotation and generation of candidate therapeutic hypotheses.

## RESULTS

### Construction of a unified lung cancer transcriptomic landscape

To construct a unified molecular reference landscape for lung cancer, we assembled transcriptomic profiles from 1,824 tumors across 17 independent studies^1,13–17^. The integrated cohort is anchored by several large publicly available lung cancer transcriptomic datasets, including TCGA-LUAD^13^ (n=529), TCGA-LUSC^1^ (n=519), lung cancer samples from CPTAC-3^18^ (n=266), APOLLO-LUAD^14^ (n=87), and 124 NSCLC samples from Bakr^19^ et al. (SRP117020). These larger cohorts were supplemented with patient tumor samples from smaller publicly available studies with clinical or histologic annotations deposited in the Gene Expression Omnibus (GEO), thereby broadening representation across lung cancer histologies and independent patient populations (Table S1).

After uniform preprocessing, batch correction and variance-stabilizing transformation (VST), we analyzed the integrated cohort using two complementary approaches. First, consensus clustering was used to define transcriptional groups from batch corrected expression values for 19,055 protein coding genes. Second, dimensionality-reduction methods were applied to the same 19,055 protein-coding gene expression data to visualize relationships among samples across cohorts and disease types.

Consensus clustering was performed using batch-corrected, variance-stabilized expression values for all protein-coding genes (Figure S1A). Because the integrated cohort contained three major histologic disease classes, we initially expected the full-cohort clustering to resolve adenocarcinoma-, squamous-, and small-cell-dominant compartments. Instead, full-cohort consensus clustering resolved three major transcriptional compartments consisting of a predominantly adenocarcinoma region, a squamous-enriched region, and a second adenocarcinoma-associated region that also contained a substantial fraction of small cell lung cancer and NSCLC(NOS) samples. This adenocarcinoma–small-cell mixed compartment was internally heterogeneous and was therefore subjected to secondary consensus clustering, which resolved five transcriptional groups designated A2, A3, A4, SC, and MD1. Separately, the squamous-enriched compartment was clustered and resolved into two squamous transcriptional groups, SQ1 and SQ2. Thus, the final landscape annotation was derived from coarse full-cohort clustering followed by targeted refinement of the compartments in which major disease-associated transcriptional structure remained unresolved.

To evaluate whether the visual organization of the clusters depended on the dimensionality-reduction method, we compared PCA, UMAP, t-SNE, and PaCMAP using identical batch-corrected protein-coding gene expression inputs. Each representation was annotated by dataset, disease, and clusters defined by consensus clustering (Figure S1B). Major disease-and cluster-level relationships were recoverable across methods, indicating that the principal organization of the cohort was not unique to PaCMAP. PaCMAP provided the clearest visual representation of both local sample neighborhoods and broader cross-histology relationships and was therefore retained as the primary visualization framework.

The PaCMAP visualization of all 1,824 tumors showed substantial mixing of samples from different studies, consistent with reduced study-specific technical structure after batch correction (Figure 1A). Annotation by pathologist-assigned diagnosis revealed two broad regions of transcriptional organization. One region was enriched for adenocarcinoma, with interspersed small cell lung cancer and NSCLC/NOS samples. The second region was enriched for squamous cell carcinoma but also contained subsets of tumors annotated as adenocarcinoma, small cell lung cancer, or NSCLC/NOS. Thus, histology provided the dominant organizing axis of the map, but did not fully explain the transcriptional structure of the integrated cohort (Figure 1B-G).

**Figure 1.**
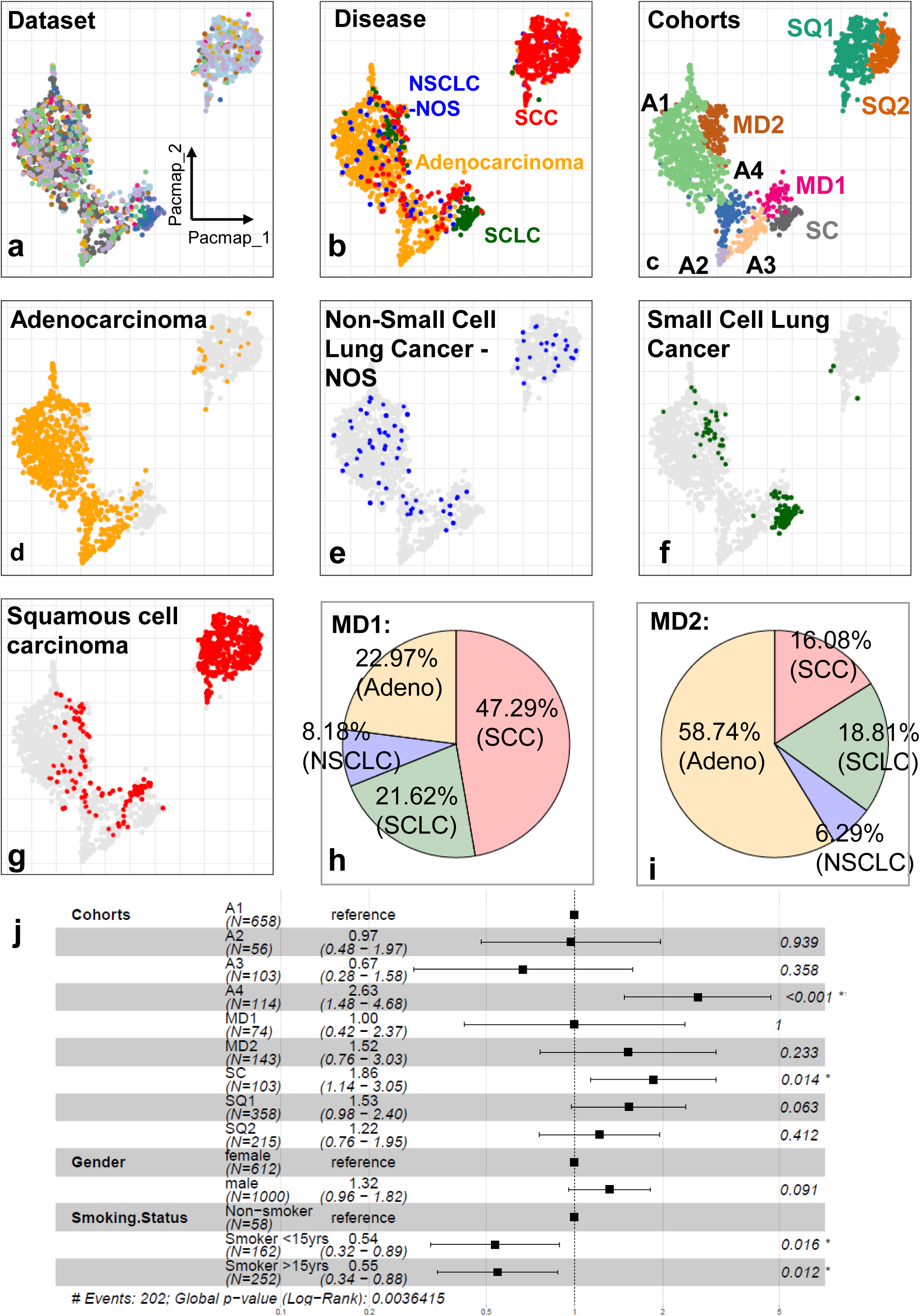
Construction of a unified lung cancer transcriptomic landscape and cohort-level clinical associations. (A) PaCMAP embedding of 1,824 lung cancer samples colored by dataset of origin following uniform processing and batch correction. Effective intermixing of datasets indicates mitigation of study-specific effects. (B) Same embedding colored by histological diagnosis, including adenocarcinoma, squamous cell carcinoma (SCC), small cell lung cancer (SCLC), and non–small cell lung cancer not otherwise specified (NSCLC-NOS). (C) PaCMAP embedding colored by transcriptional cohorts identified through consensus clustering, including A1–A4, SC, SQ1, SQ2, and mixed-diagnosis regions (MD1 and MD2). (D-G) PaCMAP embedding colored by tumors diagnosed as adenocarcinoma, non-small cell lung cancer, small cell lung cancer and squamous cell carcinoma samples. Panels D–G show diagnosis-specific overlays to highlight both the dominant region occupied by each histologic group and smaller dispersed subsets that are less apparent in the combined disease annotation shown in panel B. (H-I) Pie charts for mixed diagnosis region MD2 and MD1 showing distribution of adenocarcinoma, small cell lung cancer and squamous cell carcinoma samples. (J) Forest plot of multivariable Cox proportional hazards models evaluating overall survival across transcriptional cohorts (Reference group: A1), adjusted for cohort, sex, and smoking status. Hazard ratios are shown with 95% confidence intervals.

We identified two mixed-diagnosis regions. Adjacent to SC, we identified an additional mixed-diagnosis region (MD1), containing tumors diagnosed as 22.97% adenocarcinoma, 8.18% NSCLC/NOS, 21.62% small cell lung cancer, and 47.29% squamous cell carcinoma (Figure 1H). Within A1, the second mixed diagnosis region (MD2) was composed of tumors diagnosed as 58.74% adenocarcinoma, 6.29% NSCLC/NOS, 18.81% small cell lung cancer, and 16.08% squamous cell carcinoma (Figure 1I). The presence of histologically discordant cases within each region may reflect diagnostic ambiguity, combined histology, sampling effects, or neuroendocrine-associated transcriptional programs not fully captured by conventional pathology.

We further assessed the stability of the landscape by repeatedly reconstructing PaCMAP embeddings using 25%, 50%, 70%, and 90% of the full cohort (Figure S1C). The major disease-and cohort-level topology was preserved at 50%, 70%, and 90% sampling, whereas the 25% embedding was more dispersed and less structured. These results indicate that the major transcriptional relationships in the integrated landscape were robust to substantial cohort subsampling, while lower sample sizes reduced the stability and resolution of the visual structure.

Multivariable Cox proportional hazards modeling adjusting for cohort, sex, and smoking status showed that A4 and SC were associated with significantly poorer overall survival relative to A1 (A4 = HR 2.63, 95% CI 1.48–4.68, p < 0.001, SC=HR 1.86, 95% CI 1.14–3.05, p = 0.014). The overall model was significant (global log-rank p = 0.003) with moderate discriminative performance (concordance index 0.62), indicating that molecular organization within the landscape was associated with overall survival after adjustment for available covariates (Figure 1J, Figure S1E). Pairwise survival comparisons across all nine clusters using Tukey-adjusted Cox models showed that the prognostic signal was concentrated primarily in comparisons involving A4 and SC, rather than being broadly distributed across the landscape (Table S2). Within the mixed-diagnosis regions, histology-stratified survival analysis showed no significant survival differences in MD1, whereas histology remained prognostic in MD2, with SCLC cases showing poorer survival than LUAD and LUSC cases (Table S2).

### Canonical lineage markers and genomic features align with landscape organization

To validate that the unified landscape captures established lung cancer biology, we overlaid expression of canonical lineage markers and orthogonal genomic features derived from the literature. As expected, expression of the lung lineage transcription factor NKX2-1^20^ was enriched in regions dominated by adenocarcinoma samples and SCLC regions but reduced in squamous cell carcinoma clusters, consistent with lineage-specific transcriptional identity across the map (Figure 2A). By contrast, KRT5^21^ and other keratins, a marker of squamous differentiation, were upregulated in the squamous cell carcinoma region, reinforcing the molecular distinctiveness of this histological subtype (Figure 2B). As expected, small cell lung cancer regions were characterized by elevated expression of neuroendocrine regulators, including NEUROD1^4^, consistent with established SCLC transcriptional programs (Figure 2C).

**Figure 2.**
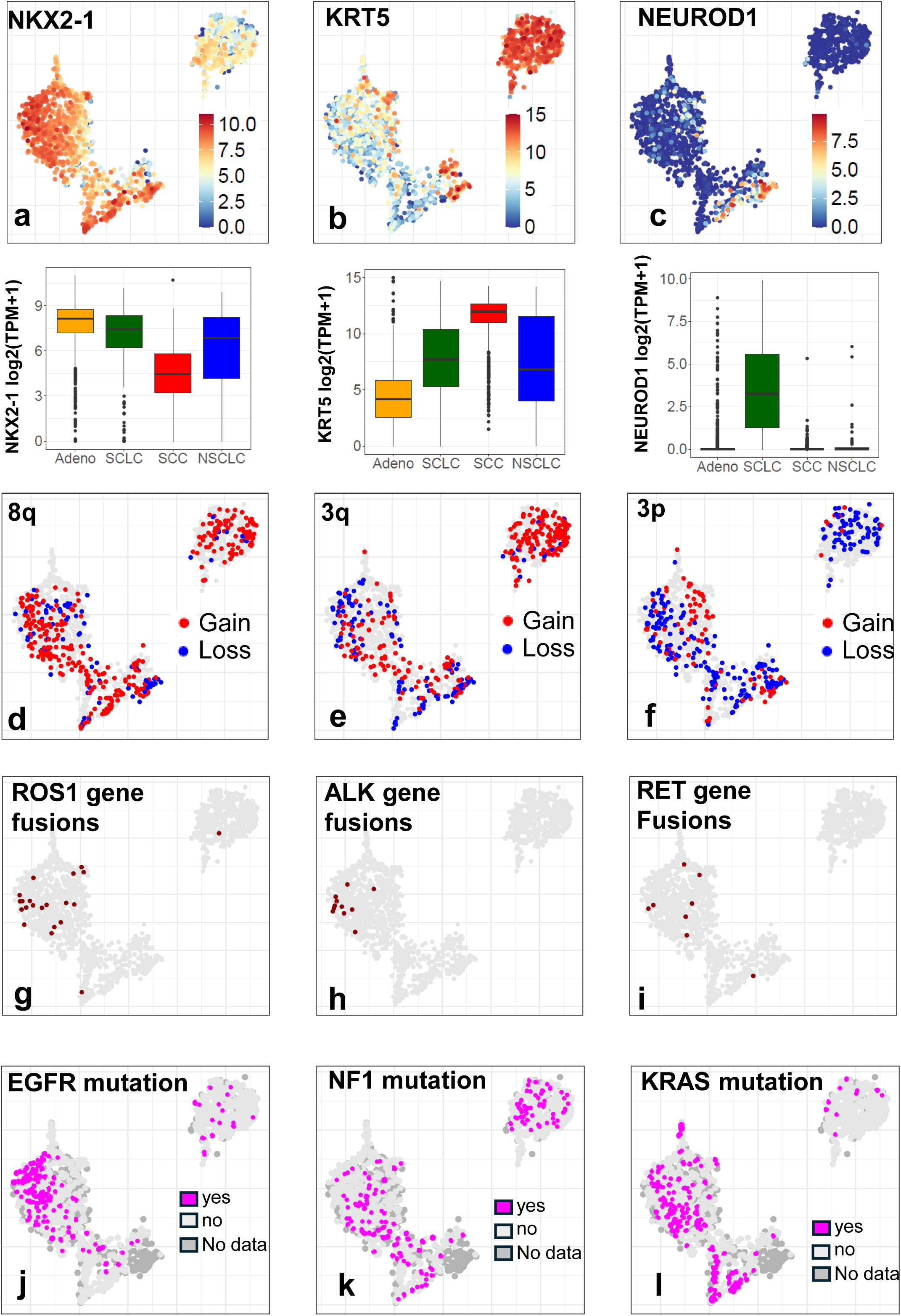
Molecular features align with spatial organization of the unified lung cancer landscape. (A–C) PaCMAP embedding colored by normalized gene expression of canonical lineage markers: NKX2-1 (adenocarcinoma-associated) (A), KRT5 (squamous lineage marker) (B), and NEUROD1 (neuroendocrine marker) (C). Gene expression intensity (log2(TPM+1)) is shown on a continuous scale. Boxplots below PaCMAP show gene expression of gene, by disease. (D–F) Inferred arm-level copy number alterations projected onto the landscape: 8q (D), 3q (E), and 3p (F), as determined from RNA-Seq–based copy number inference. Patients with amplifications or deletions in chromosome arms are shown in red and blue respectively. Patients with no copy number changes are shown in grey. (G–I) PaCMAP embedding colored by recurrent gene fusions, including ROS1, ALK, and RET. Fusion-positive samples are highlighted. (J–L) PaCMAP embedding colored by mutations - EGFR, NF1, and KRAS. Samples with mutations are highlighted.

Beyond gene expression, copy-number alteration patterns, inferred from CaSpER using bulk RNA-Seq data, mirrored known subtype-specific genomic architectures: squamous cell carcinoma were enriched for recurrent gains involving chromosomes 3q(24.95%), 5p(15.5%), 8q(15.07%), and losses on 3p(11.83%) and 8p(11.5%)^2,22^, whereas adenocarcinomas exhibited characteristic gains on 5p (17.84%), 8q (18.35%),7p (17.09%) (Figure 2D-F Table S3). NSCLC/NOS and SCLC exhibited extensive copy number aberrations (Figure S2A).

### Clinical, molecular, and tumor microenvironment annotation of transcriptional clusters

We next asked how the transcriptional landscape related to routinely available clinical and molecular annotations. Across 1,824 tumors, cluster membership was evaluated against pathologist-assigned histology, AJCC pathologic stage, smoking status, sex, oncogenotype, nonsynonymous mutation burden, neuroendocrine marker expression, and tumor microenvironment features (Figure S2D–K; Table S5).

We also annotated the landscape with recurrent oncogenic gene fusions, including ALK(1.7%), ROS1(2.3%), RET(0.8%)^13^, MET(0.8%)^23^ and NTRK2^24^ rearrangements (Figure 2G-I, Figure S2B, Table S4), inferred from Arriba. Driver-event annotations showed non-random but event-specific localization. RNA-derived fusion calls identified enrichment of ROS1 and ALK fusions in A1, NRG1 fusions in SQ1, and NTRK2 fusions in SQ2 (Table S4-S5).

Because mutation calling from RNA sequencing data is subject to important technical limitations, including confounding from RNA editing, reverse transcription artifacts, and mapping ambiguity, we leveraged matched DNA-based mutation calls available for select samples (n=1305, 71.5%) to annotate the landscape (Figure 2J-L, Figure S2C). DNA mutation calls showed strong enrichment of EGFR mutations in A1 and depletion from squamous-associated clusters, while KRAS mutations were enriched across adenocarcinoma-associated regions and depleted from SQ1 and SQ2. BRAF and MET mutations also showed significant cluster association. Nonsynonymous mutation burden differed across clusters, with higher burden in A3, A4, MD1, SQ1, and SQ2 and lower burden in A1, A2, and MD2. Because callable territory was not uniform across cohorts, mutation burden was analyzed as nonsynonymous mutation count rather than formal mutations per megabase.

Pathologist-assigned histology showed a highly significant association with cluster membership (global Fisher’s exact test with simulated p-value, FDR = 3.70 × 10□□) (Figure S2E, Table S5). A1–A4 were enriched for LUAD, with A2 composed entirely of LUAD cases and A1 containing 594 LUAD tumors among 658 cases (90.3%). SQ1 and SQ2 were strongly enriched for LUSC/SCC, comprising 323/358 cases (90.2%) and 201/215 cases (93.5%), respectively, while SC was strongly enriched for SCLC, comprising 98/103 cases (95.1%). In contrast, MD1 and MD2 showed mixed histologic composition, with significant SCLC enrichment in both regions, supporting their designation as mixed-diagnosis transcriptional regions rather than histology-specific clusters.

Clinical annotations showed additional cluster-specific structure. Stage varied significantly across clusters, with A1 enriched for stage I disease, SC enriched for stage III disease, and A4 enriched for stage IV disease (Figure S2F, Table S5). Stage IV tumors were relatively uncommon, likely reflecting the enrichment of public bulk RNA-seq cohorts for resected primary tumors and the limited availability of surgical tissue from advanced-stage disease. Despite this limitation, stage IV cases were not confined to a single transcriptional region, indicating that stage alone did not explain the global organization of the landscape.

Sex distribution also differed across the landscape (Figure S2G, Table S5), with A1 significantly enriched for female patients and SQ1, SQ2, and SC enriched for male patients. Smoking status was available for a subset of tumors and varied by cluster: A1 was enriched for non-smokers, whereas SQ1 and SQ2 were enriched for current smokers. Because smoking annotation was incomplete and pack-year exposure was not uniformly available, smoking associations were interpreted as cluster-level clinical annotations rather than quantitative exposure effects.

Tumor microenvironment and neuroendocrine annotations (Figure S2J-K, Table S5) further refined cluster interpretation. Orthogonal deconvolution approaches showed that SQ1 had features of an immune-enriched squamous state, including higher inferred CD8 T-cell and M1 macrophage fractions, while MD2 showed higher inferred monocyte and CD8 T-cell fractions, relative to other clusters. By contrast, A4 showed lower immune and microenvironment scores across multiple methods, and SQ1 showed lower CD274 expression and lower xCell immune score despite lower PUREE purity, indicating that reduced purity did not necessarily reflect a strongly inflamed state. Neuroendocrine marker expression varied strongly across clusters, with highest scores in SC and strong enrichment in A3, supporting its neuroendocrine-like adenocarcinoma-associated phenotype.

Specimen type was also evaluated where metadata were available; all tumors with available specimen annotation in the analyzed cohort were annotated as primary tumors, and metastatic behavior could not be further assessed from the available metadata. Together, these analyses show that landscape position is shaped primarily by histology and lineage-associated transcriptional programs, with clinical variables, driver alterations, mutation burden, and tumor microenvironment features providing additional cluster-specific annotation rather than fully explaining the global organization of the atlas.

### Relationship of the transcriptional landscape to established lung cancer molecular subtypes

To determine how the transcriptional landscape relates to established histology-specific molecular classifications, we directly compared the nine landscape clusters with previously defined LUAD, LUSC, and SCLC expression subtypes (Figure 3, Table S6). For LUAD and LUSC, available published subtype annotations were projected onto the landscape and compared side-by-side with subtype assignments generated using published subtype-associated gene signatures. For SCLC, samples were classified using the SCLC-A, SCLC-N, SCLC-P, and SCLC-I framework described by Gay et al., with low-margin assignments retained as ambiguous. Because the goal was to evaluate how disease-specific subtype programs are represented within the cross-histology landscape clusters, subtype percentages were calculated across all samples in each landscape cluster; samples without an applicable inferred subtype assignment are shown as “Not Classified,” and samples lacking published subtype annotation are shown as “Not Reported.”

**Figure 3.**
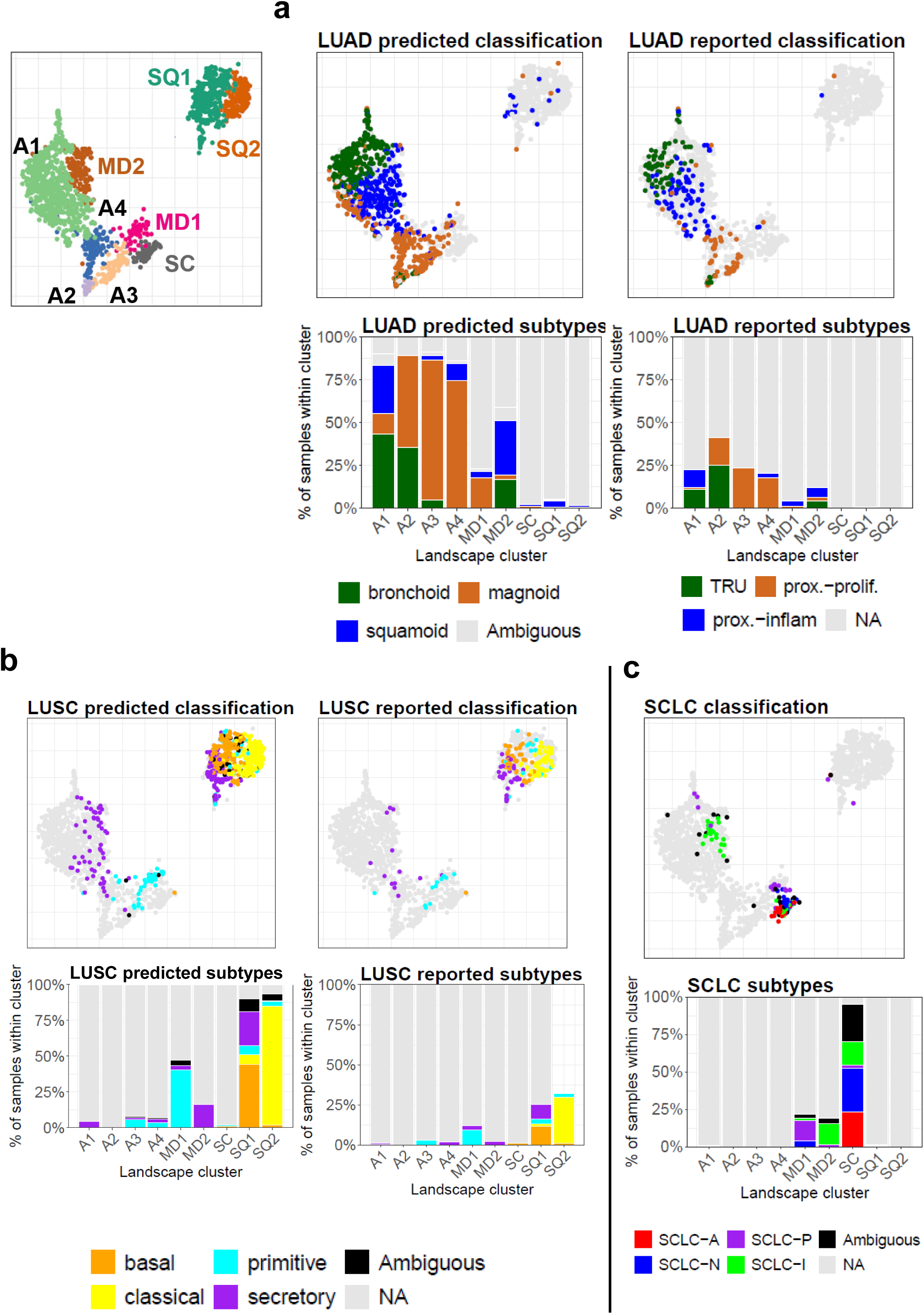
Relationship between landscape clusters and established lung cancer molecular subtype frameworks. (A) LUAD subtype annotations projected onto the integrated PaCMAP landscape. The upper panels show samples colored by inferred LUAD subtype labels and reported LUAD subtype labels when available. The lower stacked barplots show the distribution of LUAD subtype assignments within each transcriptional landscape cluster. Inferred LUAD subtypes were assigned using published subtype-associated gene programs, while reported subtype labels were retained from the original source studies. Samples without an applicable inferred LUAD subtype assignment are labeled “Not Classified,” and samples lacking reported subtype annotation are labeled “Not Reported.” (B) LUSC subtype annotations projected onto the integrated PaCMAP landscape. The upper panels show samples colored by inferred and reported LUSC subtype labels, and the lower stacked barplots show subtype composition within each transcriptional landscape cluster. Inferred LUSC subtypes were assigned using published basal, classical, primitive, and secretory subtype programs. “Not Classified” denotes samples without an applicable inferred LUSC subtype assignment, and “Not Reported” denotes samples without reported subtype annotation. (C) SCLC subtype annotations projected onto the integrated PaCMAP landscape. The upper panel shows samples colored by inferred SCLC subtype assignment according to the SCLC-A, SCLC-N, SCLC-P, and SCLC-I framework, with low-confidence assignments labeled ambiguous. The lower stacked barplot shows the distribution of inferred SCLC subtype assignments across transcriptional landscape clusters.

For LUAD, inferred bronchoid/TRU, magnoid/proximal-proliferative, and squamoid/proximal-inflammatory classes showed broad spatial concordance with available published annotations (Figure 3A). A1 contained a mixture of inferred bronchoid (43.2%), squamoid (28.4%), and magnoid (11.9%) assignments, consistent with the presence of both TRU-and proximal-inflammatory-associated tumors among the available reported annotations in this region. In contrast, A2, A3, and A4 were enriched for the magnoid program, comprising 53.6%, 81.6%, and 74.6% of each cluster, respectively. Available reported annotations showed a concordant pattern, with proximal-proliferative labels concentrated in A2–A4 and comparatively rare in A1.

A similar pattern was observed for the basal, classical, primitive, and secretory LUSC expression subtypes (Figure 3B). SQ2 was dominated by the classical program, with 83.3% of samples classified as classical and a concordant predominance of classical labels among the available reported annotations. In contrast, SQ1 was more heterogeneous, containing basal (44.4%), secretory (23.7%), classical (6.7%), and primitive (6.2%) inferred assignments, with 9.2% ambiguous assignments. Established LUSC programs also appeared outside the principal squamous clusters: MD1 contained a prominent primitive component (40.5%), while MD2 contained a smaller secretory component (16.1%).

SCLC subtype programs were also distributed non-uniformly across the landscape (Figure 3C). The SC cluster contained the largest concentration of classified SCLC samples included SCLC-N (29.1%), SCLC-A (23.3%), SCLC-I (15.5%), and SCLC-P (1.9%) assignments, along with a substantial ambiguous fraction (25.2%). In addition, SCLC-P assignments were enriched in MD1 (13.5%), whereas SCLC-I assignments were enriched in MD2 (14.0%).

To formally quantify the relationship between the nine landscape clusters and established molecular subtype systems, we performed cluster-by-subtype concordance analyses using disease-matched samples with available reported or inferred subtype assignments (Table S6). Landscape clusters were significantly associated with prior subtype labels for LUAD, LUSC, and SCLC (Monte Carlo chi-square tests, all p < 0.001). The strength of association was moderate to strong across frameworks, with Cramer’s V values of 0.516 and 0.593 for predicted and reported LUAD subtypes, 0.692 and 0.622 for predicted and reported LUSC subtypes, and 0.539 for predicted SCLC subtypes. However, adjusted Rand index and normalized mutual information values showed only partial agreement between the nine landscape clusters and prior subtype systems, indicating that the clusters recover established subtype-associated biological programs but are not reducible to existing discrete classifications.

### Adenocarcinoma-associated and mixed-diagnosis regions show distinct transcriptional programs

Differential gene expression analysis revealed that A2–A4, MD1 and MD2 were characterized by distinct and non-overlapping biological programs relative to A1, consistent with the presence of multiple tumor-intrinsic transcriptional states (Figure 4A).

**Figure 4.**
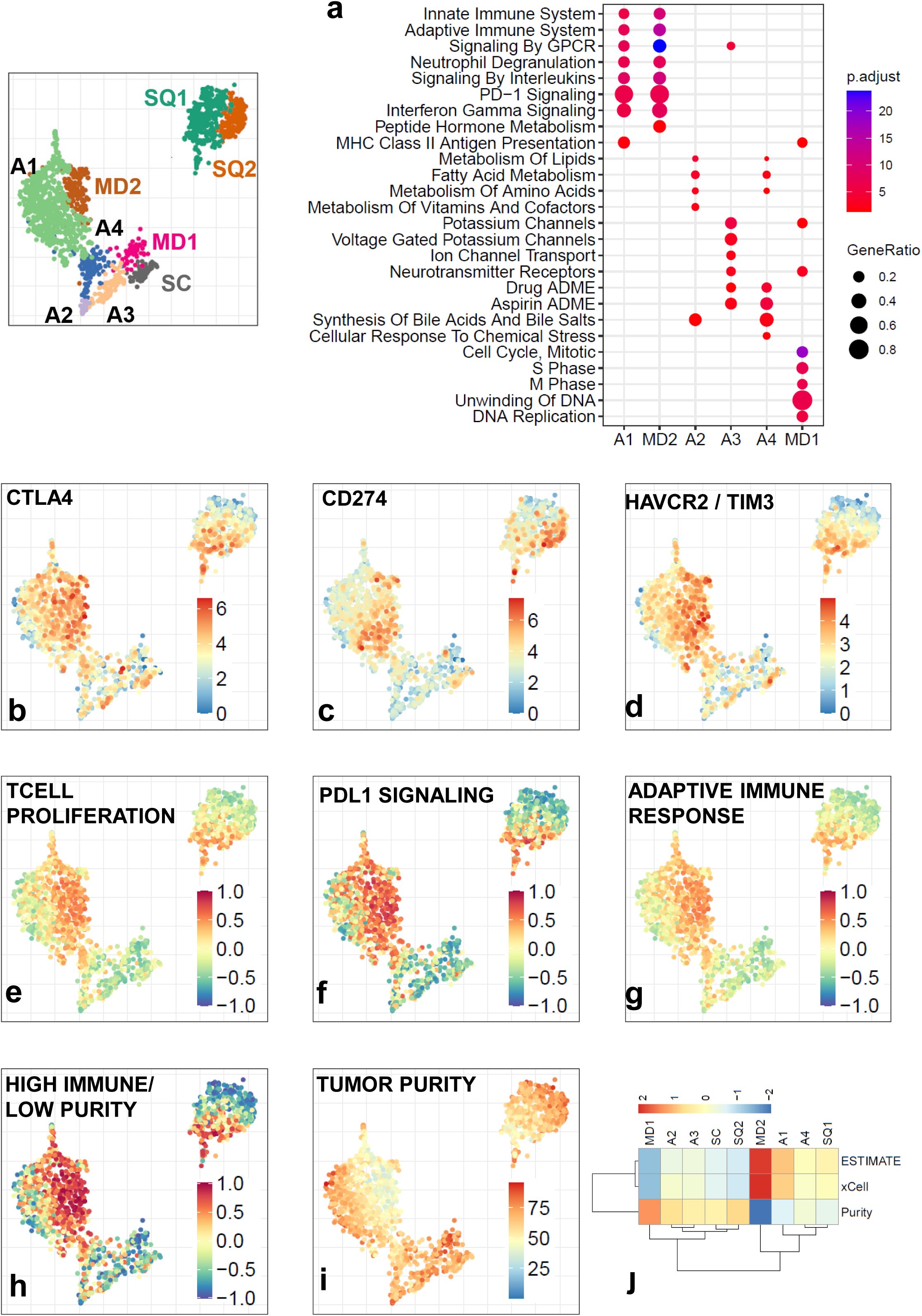
Adenocarcinoma-associated and mixed-diagnosis regions show distinct transcriptional programs. (A) Dot plot summarizing pathway enrichment across A1–A4, MD1 and MD2, based on GSVA scores. Dot size reflects the proportion of samples with elevated pathway activity, and color intensity indicates relative pathway enrichment. (B-D) PaCMAP embedding colored by normalized gene expression of immune-related markers: CTLA4 (B), CD274 (PD-L1) (C) and TIM3(D). Expression intensity is shown on a continuous scale. (E-G) PaCMAP embedding colored by GSVA pathway activity scores for T cell proliferation (E), PD-1 signaling (F), and adaptive immune response (G). (H) PaCMAP embedding colored by GSVA scores derived from a five-gene immune predictor signature. (I) PaCMAP embedding colored by estimated tumor purity. Tumor purity was inferred using PUREE. (J) heatmap showing median score for tumor fraction estimate and tumor purity for each cluster (ESTIMATE, xcell and PUREE)

Tumors within **A2** were distinguished by enrichment of lipid and fatty acid metabolism pathways, including upregulation of FASN, alongside activation of oxidative stress response components within the NRF2 axis, including NQO1. Enrichment of NFE2L2-mediated nuclear events further indicated oxidative stress adaptation and metabolic rewiring. (Table S7-8)

**A3** consisted with tumors diagnosed as adenocarcinoma but exhibited a pronounced neuroendocrine transcriptional program, with robust upregulation of canonical lineage regulators including ASCL1, NEUROD1, INSM1, CHGA, CHGB, DLL3, MYCN, MYCL, RET, SYP, and NCAM1 (Figure S3, Table S7). In total, 1,090 genes were significantly upregulated in A3 relative to A1. The coordinated induction of master neuroendocrine transcription factors together with synaptic vesicle machinery genes indicates activation of a lineage-defining neuroendocrine state rather than isolated pathway enrichment. Notably, YAP1 and POU2F3 were not enriched in A3, suggesting that this subgroup does not correspond to the SCLC-Y or SCLC-P transcriptional states. The absence of significant NKX2-1 differential expression further supports preservation of lung lineage identity within this neuroendocrine-like adenocarcinoma context. Together, these findings define A3 as a neuroendocrine-like adenocarcinoma state consistent with lineage plasticity across traditional histological boundaries (Table S7-8).

Tumors within **A4** demonstrated strong enrichment for xenobiotic metabolism pathways, with significant upregulation of multiple cytochrome P450 enzymes (CYP24A1 (Figure S4A), CYP2C18, CYP2C19, CYP2C9, CYP4F11, CYP4F2, CYP4F3), consistent with enhanced carcinogen and drug metabolism.

This was accompanied by activation of glutathione synthesis and redox defense genes (GCLC (Figure S4A), GSR, GSTP1) and ferroptosis-associated regulators, including SLC7A11(Figure S4A), indicating increased oxidative stress adaptation. Concurrent upregulation of pentose phosphate pathway enzymes (G6PD, PGD) further suggests metabolic rerouting toward NADPH generation and antioxidant buffering. Collectively, these features define A4 as a detoxification-competent, redox-adapted adenocarcinoma state, potentially reflecting transcriptional remodeling in response to environmental stressors. (Figure 3B, Figure S3, Table S7-8)

The diagnosis-discordant region (MD1) was characterized by coordinated upregulation of core cell cycle regulators including CDK1, CDK2, and E2F1 (Figure S4A), consistent with active G1/S and G2/M transition control. This proliferative signature was reinforced by robust induction of DNA replication machinery components (MCM2–7, PCNA, POLA1) and mismatch repair genes (MSH2, MSH6). Enrichment of mitotic checkpoint regulators (BUB1, AURKA, PLK1) further supports a state of heightened mitotic activity and checkpoint engagement. Together, these findings define MD1 as a mixed-diagnosis transcriptional region marked by proliferative acceleration and potential replication stress, consistent with a genomically unstable phenotype (Table S7-8).

### An immune-infiltrated adenocarcinoma state exhibits coordinated adaptive and innate immune activation

We next examined the global transcriptional programs characterizing A1 and MD2. Gene expression analysis revealed strong enrichment of adaptive immune markers, including CD3D, CD3E, CD4, CD8A, CD8B, LCK, and ZAP70, indicating increased T cell infiltration within A1 and MD2. Immune checkpoint molecules were also significantly upregulated, including PDCD1 (PD-1), CD274 (PD-L1), CTLA4, LAG3, and TIGIT, consistent with an activated yet potentially exhausted immune microenvironment (Figure 4A-G).

Interferon signaling pathways were prominently induced in A1 and MD2, with elevated expression of STAT1, CXCL9, CXCL10, and GBP1, supporting a coordinated interferon-γ response axis. In parallel, myeloid and neutrophil-associated genes including MPO, S100A8, and CXCR2 were significantly enriched, indicating engagement of innate immune components. Finally, extracellular matrix and stromal remodeling genes such as COL1A1, COL3A1, MMP9, and FN1 were upregulated, consistent with active tumor–stroma interactions and immune cell trafficking within this region (Table S7-8).

To complement the immune-gene expression analysis in A1 and MD2, we examined tumor purity as an orthogonal indicator of non-tumor cellular contribution. A1 and MD2 showed lower PUREE-estimated tumor purity relative to the more tumor-intrinsic adenocarcinoma-associated states (Figure 4H–J). In parallel, a GSVA score based on genes previously reported to be negatively correlated with tumor purity in lung cancer, including CD4, SASH3, CD53, PLEK, and EVI2B, was elevated in A1 and MD2 (Figure. 4H, Figure S4B). These findings are concordant with prior reports^25,26^ that support the interpretation that the immune-enriched transcriptional profile of A1 and MD2 reflects increased immune/lymphoid admixture rather than purely tumor-intrinsic expression differences. This targeted purity analysis is consistent with the broader tumor microenvironment annotation across all clusters, which showed that immune and stromal features are structured across the landscape and are not fully captured by tumor purity alone (Figure S2J, Figure 4H-J, Figure S4C).

### Small cell lung cancer segregates into proliferative and YAP1-associated immune states

Small cell lung cancer is organized into transcriptionally distinct states that include classical neuroendocrine subtypes (SCLC-A and SCLC-N) and a non-neuroendocrine YAP1-driven subtype (SCLC-Y)^4^. In the unified landscape, SCLC samples localized in two spatial domains (Figure 5A): a discrete SCLC-dominant cluster (SC) and a mixed-diagnosis region embedded within the immune-enriched adenocarcinoma domain (MD2/A1).

**Figure 5.**
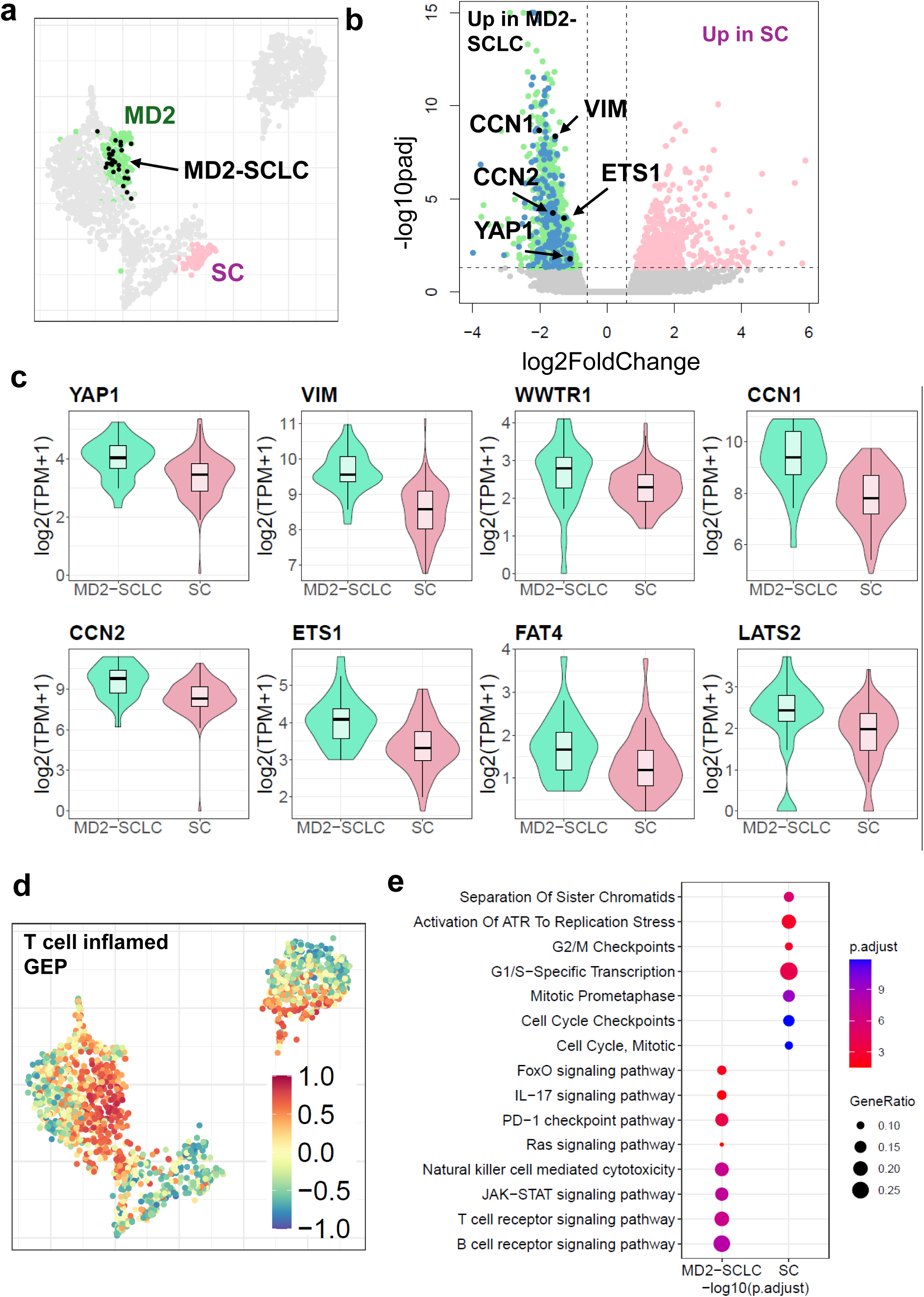
Small cell lung cancer segregates into proliferative and YAP1-associated immune states. (A) PaCMAP embedding highlighting the discrete SCLC cluster (SC) and SCLC tumors localized within the mixed-diagnosis region MD2. (B) Volcano plot of differential gene expression comparing SCLC tumors in SC versus SCLC tumors in MD2. Selected genes associated with YAP1 signaling and mesenchymal programs (CCN1, CCN2, YAP1, VIM, and ETS1) are highlighted. (C) Violin plots showing expression levels of YAP1, VIM, WWTR1, CCN1, CCN2, ETS1, FAT4, and LATS2 in SC and SCLC tumors within MD2. Higher scores are observed in SCLC tumors within MD2. (D) PaCMAP embedding colored by GSVA scores derived from the 18-gene T cell–inflamed signature. (E) Pathway enrichment analysis comparing SC and SCLC tumors in MD2, highlighting differential activation of proliferative and immune-associated pathways. Dot size represents the proportion of samples with elevated pathway activity; color intensity reflects relative enrichment.

To define the molecular programs distinguishing these regions, we performed differential gene expression analysis comparing SCLC-diagnosed samples in SC to SCLC-diagnosed tumors within MD2 (Figure 5A,B). The discrete SCLC cluster (SC) retained enrichment of canonical neuroendocrine regulators, including ASCL1, NEUROD1, and INSM1, consistent with a classical neuroendocrine phenotype. By contrast, SCLC tumors localized within MD2 exhibited significant upregulation of YAP1, VIM and WWTR1, indicating alignment with the SCLC-Y subtype (Figure 5C, Table S7).

Tumors diagnosed as SCLC and falling into MD2 also demonstrated relative enrichment of immune-associated pathways, including cytokine signaling and inflammatory responses, mirroring the transcriptional features observed in the immune-infiltrated A1 adenocarcinoma state, relative to SCLC-diagnosed tumors in SC cluster. The co-localization of MD2 with the A1 domain suggests that a subset of tumors diagnosed as SCLC reside within an immune-active microenvironment. Consistent with this interpretation, the SCLC-Y subtype has been reported to exhibit a T-cell–inflamed phenotype characterized by elevated scores on the 18-gene T-cell–inflamed expression signature (Figure 5D, Table S9)^25,27^.

By contrast, the tumors diagnosed as SCLC from the SC cluster (Figure 5A) were characterized by strong enrichment of proliferative and mitotic pathways, including cell cycle progression, mitotic spindle assembly, sister chromatid cohesion, and G2/M checkpoint regulation relative to the SCLC diagnosed tumors in MD2. Reactome pathways such as Cell Cycle, Mitotic, Mitotic Prometaphase, Polo-like kinase–mediated events, ATR activation in response to replication stress, and G1/S transcriptional regulation were significantly enriched, consistent with a highly proliferative tumor-intrinsic state (Figure 5E, Figure S5). The proximity of SC to the replication-driven adenocarcinoma cluster (A4, MD1) further supports shared proliferative transcriptional programs across histological boundaries.

Together, these findings suggest that tumors diagnosed as SCLC partitioned into at least two biologically distinct states. The first is a highly proliferative, replication-stress–associated cluster (SC) and second is an immune-associated region of tumors that are given several diagnoses (MD2) embedded within the immune-inflamed region of the map.

### Squamous cell carcinoma segregates into redox-adapted and immune-infiltrated states

Consensus clustering of squamous cell carcinoma samples onto the unified landscape revealed two spatially distinct domains – SQ1 and SQ2 (Figure 6A, Figure S1A). Differential expression analysis demonstrated that SQ1 exhibited robust activation of adaptive immune pathways, including elevated expression of CD3D, CD3E, CXCL9, CXCL10, and PDCD1, consistent with T cell infiltration and inflammatory signaling (Figure 6B-C, Figure 4). Additional enrichment of cytokine–receptor interactions, JAK–STAT signaling and processes associated with vascular remodeling and histone citrullination (Fig 6SA, Table S7,S10) further supports SQ1 as an immune-inflamed squamous carcinoma state.

**Figure 6.**
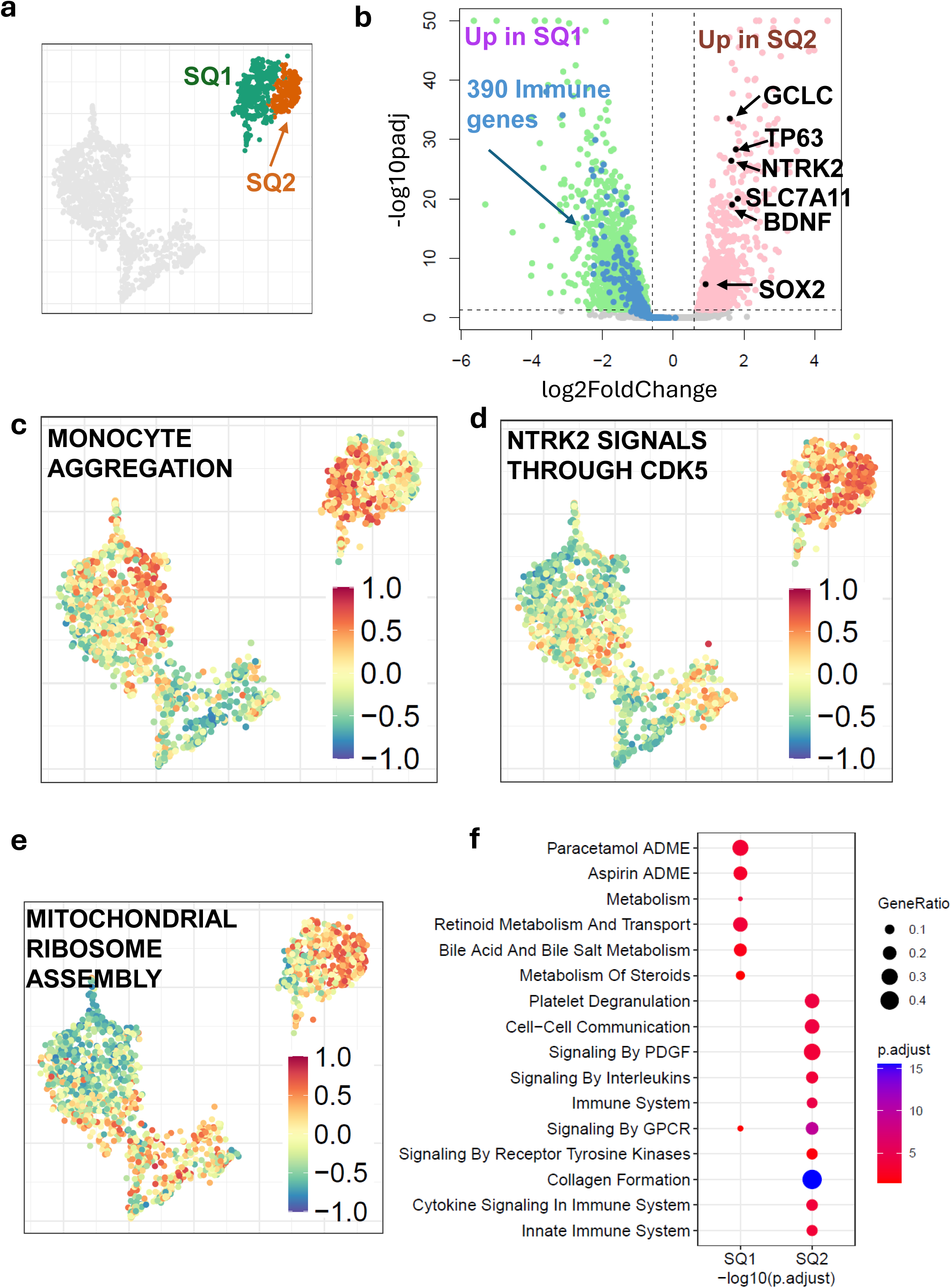
Squamous carcinoma segregates into redox-adapted and immune-infiltrated states. (A) PaCMAP embedding highlights squamous carcinoma clusters SQ1 and SQ2. (B) Volcano plot of differential gene expressions comparing SQ1 and SQ2. NTRK2 and BDNF are highlighted as upregulated in SQ2, while representative immune-associated genes upregulated in SQ1 are indicated. (C-E) PaCMAP embedding colored by GSVA pathway activity scores for monocyte, enriched in SQ1, NTRK2 signaling and ribosomal assembly enriched in SQ2 (F) Dot plot summarizing pathway enrichment differences between SQ1 and SQ2, with dot size representing the proportion of samples with elevated pathway activity and color intensity reflecting relative enrichment.

In contrast to the immune-infiltrated SQ1 state, SQ2 demonstrated selective upregulation of NTRK2 (TrkB) (logFC 1.40, padj 2.3×10□□, Figure 6C-D) along with its cognate ligand BDNF (logFC 1.15, padj 2.1×10□□), suggesting activation of a TrkB-associated signaling axis within this squamous carcinoma subset. This enrichment was specific to NTRK2, as NTRK1 and NTRK3 were preferentially expressed in SQ2. Notably, SQ2 was also enriched for NTRK2 gene fusions (Supp Fig 2, Table S5). Beyond receptor signaling, SQ2 demonstrated coordinated activation of redox and metabolic programs, including elevated SLC7A11 and GCLC expression, alongside enrichment of calcium transport, quinone metabolism (phylloquinone and menaquinone), leukotriene metabolism, mitochondrial ribosome assembly, and RNA modification pathways (Figure 6C,E-F; Figure S6A). Canonical squamous lineage markers SOX2 and TP63 were maintained, supporting lineage stability despite metabolic adaptation.

Together, these findings indicate that squamous carcinoma partitions into an immune-infiltrated state and a metabolically adapted, redox-buffered tumor-intrinsic state within the unified landscape.

### Gene expression profiles nominate region-specific candidate therapeutic hypotheses

Having defined lineage and microenvironmental states across lung cancer subtypes, we next examined whether selected clinically relevant or potentially actionable transcripts showed spatially restricted expression within the integrated landscape. Overlaying expression of targetable oncogenes revealed distinct regional enrichment patterns (Figure 7A).

**Figure 7.**
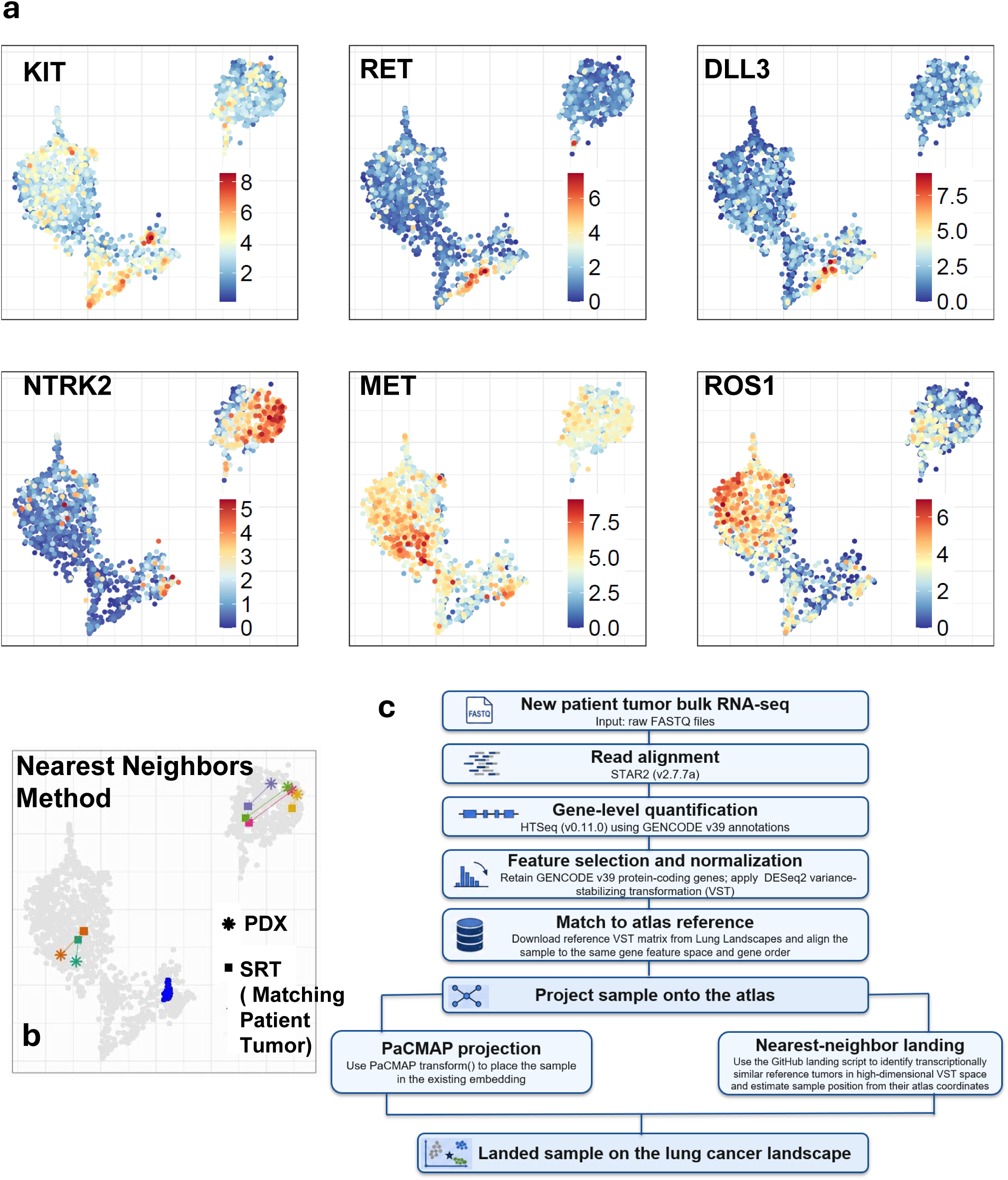
Spatially restricted potential therapeutic targets and validation of model fidelity within the unified landscape. (A) PaCMAP embedding colored by normalized gene expression of selected clinically relevant and potentially actionable transcripts, including KIT, RET, DLL3, NTRK2, MET, and ROS1. Expression intensity is shown on a continuous scale, illustrating spatial restriction of oncogenic programs across transcriptional states. These expression patterns are intended to nominate hypotheses for future functional and clinical validation and should not be interpreted as evidence of therapeutic response or as treatment-selection criteria. (B) Projection of external patient-derived xenograft (PDX) models and matched surgically resected tumors (SRTs) onto the reference landscape. Adenocarcinoma, squamous carcinoma, and small cell lung cancer PDX samples localize to their corresponding tumor-dominated regions, demonstrating preservation of transcriptional identity across independent cohorts. (C) Workflow for projecting new patient tumors or patient-derived xenograft models onto the reference lung cancer transcriptomic landscape.

KIT and RET expression was enriched in discrete regions corresponding to neuroendocrine-associated states, including the A3 adenocarcinoma cluster and portions of small cell lung cancer. Similarly, DLL3, a lineage marker and therapeutic target in small cell lung cancer, was enriched in neuroendocrine-dominant regions of the map. By contrast, NTRK2 expression was concentrated within the metabolically adapted squamous carcinoma state (SQ2), consistent with activation of a TrkB-associated tumor-intrinsic program. MET expression was elevated in specific adenocarcinoma domains, while ROS1 expression localized to restricted regions within the adenocarcinoma landscape.

These patterns were not uniformly distributed across histological categories but instead aligned with transcriptionally defined regions of the landscape. Thus, the integrated map provides a framework for nominating region-specific therapeutic hypotheses. Functional testing and validation in clinically annotated cohorts with treatment-response data will be required to determine whether any of these RNA-defined regions correspond to actionable therapeutic vulnerabilities.

### Projection of patient-derived xenografts onto the reference landscape

To assess how well the unified landscape is able to contextualize experimental models, we projected independent patient-derived xenograft (PDX) transcriptomes from two published studies^28,29^ onto the reference embedding (Figure 7B, Table S11), including 43 small cell, 4 squamous, and 2 adenocarcinoma PDX models, using two independent pipelines (Figure 7B, Figure S7B). Across histologies, PDX samples localized to the corresponding tumor-dominated regions of the landscape: adenocarcinoma PDXs mapped within adenocarcinoma states, small cell PDXs aligned with small cell–associated regions, and squamous PDXs clustered within squamous carcinoma domains. These projections recapitulated the transcriptional identity of the originating tumor types, indicating that the atlas preserves biologically meaningful structure across independent cohorts.

For adenocarcinoma and squamous carcinoma, where matched initial surgically resected tumors (SRTs; n = 6) and their corresponding PDX models were available, direct comparison revealed close spatial proximity between each patient tumor and its derived xenograft (Figure 7B). In several cases, SRT–PDX pairs occupied nearly overlapping coordinates within the embedding, supporting preservation of tumor-intrinsic transcriptional programs during engraftment and reinforcing the utility of the landscape as a quantitative framework for model validation.

A workflow for projecting individual patient tumors or patient-derived xenografts onto the reference landscape is illustrated in Figure 7C and described in detail in the Methods. Together, these findings demonstrate that the unified landscape not only resolves endogenous tumor heterogeneity but also provides a quantitative framework for assessing model fidelity and determining how accurately experimental systems recapitulate patient tumor states.

## DISCUSSION

Lung cancer has historically been classified according to histopathological criteria that define major subtypes, including adenocarcinoma, squamous cell carcinoma, and small cell lung cancer. While this framework has guided clinical management, it incompletely captures the molecular and biological heterogeneity that exists both within and across these categories. By integrating 1,824 tumors into a unified transcriptomic landscape, we show that bulk RNA-seq profiles from lung cancer can be organized as a structured continuum of transcriptional states that complements conventional histopathologic classification.

The contribution of this study is threefold. Conceptually, the landscape provides a shared cross-histology transcriptomic framework in which established LUAD, LUSC, and SCLC subtype programs can be compared within the same reference space, rather than as isolated disease-specific classifications. Methodologically, the study integrates uniformly processed bulk RNA-seq data across multiple independent cohorts and derives transcriptional structure from genome-scale protein-coding gene expression rather than from predefined subtype signatures alone. Operationally, the resulting resource is deployed as an interactive reference that allows users to examine tumors, clusters, molecular annotations, copy-number alterations, fusions, immune features, and candidate programs within a common lung cancer landscape. Thus, the atlas is best viewed not as replacing existing histologic or molecular subtype systems, but as organizing them within a broader reference framework.

Mapping routine clinical and molecular annotations onto the landscape showed that some variables tracked strongly with transcriptional position, whereas others provided contextual annotation without dominating the global structure. Pathologist-assigned histology was the strongest clinical correlate, with LUAD-, LUSC/SCC-, and SCLC-enriched regions occupying distinct areas of the map, while MD1 and MD2 captured mixed-diagnosis transcriptional neighborhoods. Neuroendocrine marker expression and tumor microenvironment features also showed strong regional structure, distinguishing SC and A3 as neuroendocrine-associated regions and identifying immune-or stromal-enriched compartments such as SQ1 and MD2. By contrast, sex, smoking status, stage, mutation burden, and individual oncogenic alterations showed cluster-specific enrichments but did not fully explain the global organization of the landscape. These results suggest that landscape position is primarily shaped by histology, lineage state, and microenvironmental programs, with routine clinical and molecular variables contributing additional but incomplete explanatory structure.

Across histologies, the unified landscape revealed that lung cancer is organized along two conserved axes: tumor-intrinsic programs defined by proliferative or metabolic adaptation, and immune-infiltrated states marked by coordinated adaptive and innate immune activation. Tumors diagnosed as adenocarcinoma largely resolved into five transcriptional states, including metabolically rewired, neuroendocrine-like, xenobiotic/redox-adapted, proliferative, and immune-enriched subtypes. The neuroendocrine-like adenocarcinoma cluster characterized by ASCL1, NEUROD1, and DLL3 expression raises the possibility of neuroendocrine-associated transcriptional convergence across conventional diagnostic boundaries. Tumors diagnosed as small cell lung cancer followed a parallel architecture, segregating into a proliferative neuroendocrine state (SC) and a YAP1-associated immune state (MD2), while squamous carcinoma partitioned into a redox-adapted, TrkB-enriched tumor-intrinsic state (SQ2) and an immune-infiltrated state (SQ1). The recurrence of this architecture across diseases suggests that transcriptional state may complement histological origin in defining tumor biology.

Immune-enriched states across lung cancer histologies shared upregulation of T cell markers, interferon signaling, and checkpoint molecules and were associated with reduced tumor purity, consistent with an inflamed tumor microenvironment. Conversely, tumor-intrinsic states displayed coordinated metabolic and survival adaptations: proliferative adenocarcinoma and small cell clusters converged on replication stress and cell cycle programs, whereas the redox-adapted squamous state exhibited enhanced glutathione biosynthesis and pentose phosphate activity alongside NTRK2 and BDNF expression. Together, these findings suggest that lineage identity intersects with conserved metabolic and immune axes to define stable yet context-dependent tumor states.

The landscape may also provide a roadmap for functional follow-up by identifying the transcriptional contexts in which candidate target programs are enriched. The spatial restriction of potentially actionable transcripts, including KIT, RET, DLL3, NTRK2, MET, and ROS1, suggests that the landscape may help nominate therapeutic hypotheses for future study. However, these hypotheses are based on bulk RNA expression and should not be interpreted as evidence of functional dependency or drug sensitivity. A practical next step is to test cluster-associated candidate programs using perturbational screens, pharmacologic assays, organoid or PDX models, immune co-culture systems, and treatment-annotated clinical cohorts. We therefore provide a candidate functional-testing framework to guide future studies rather than to define validated therapeutic vulnerabilities (Table S12).

Notably, the presence of mixed-diagnosis regions (MD1 and MD2) highlights transcriptional neighborhoods with heterogeneous pathological diagnoses and shared expression structure. Importantly, survival analyses demonstrate that outcomes of these transcriptionally defined subregions can differ, suggesting that landscape position may help generate hypotheses for future cohort stratification. In prospective use, RNA-seq placement of a tumor within an MD region may prompt pathology re-review.

Investigators can project compatible patient transcriptomes onto the reference map to refine molecular subdiagnosis, contextualize ambiguous cases, compare tumors with nearby molecularly annotated regions, and generate hypotheses for prospective studies. Proximity to well-characterized landscape regions may nominate testable hypotheses regarding tumor biology and candidate targetable programs.

Similarly, experimental models, including patient-derived xenografts, can be projected onto the landscape to assess model fidelity and determine how closely they recapitulate specific tumor states. Our observation that matched surgically-resected tumors and their matched PDX derivatives occupy closely aligned coordinates supports preservation of transcriptional identity during engraftment.

To facilitate broad access, the atlas is integrated with an interactive web interface using Oncoscape, enabling investigators to iteratively query genes or pathways of interest and visualize their spatial distribution across lung cancer states. This resource allows real-time exploration of transcriptional programs and provides a community-accessible framework for hypothesis generation and comparative analysis.

In summary, this unified lung cancer landscape shows that histological categories intersect with conserved tumor-intrinsic and immune axes of organization. By organizing lineage-associated programs, microenvironmental engagement, and spatially structured candidate target programs within a single reference framework, this atlas provides a scalable resource for biological discovery, model evaluation, and hypothesis generation across lung cancer.

### Limitations of the study

Several caveats warrant consideration. First, a major limitation of the current landscape is that each tumor is represented by a single bulk RNA-seq profile and therefore by a single coordinate in the reference space. This representation collapses intratumor transcriptional heterogeneity and may obscure spatially distinct compartments, such as immune-infiltrated peripheral regions, proliferative tumor cores, or lineage-divergent subclones. Consequently, landscape position should be interpreted as a composite representation of the sampled bulk tissue rather than as a complete description of all transcriptional states within an individual tumor. Future studies using multiregion bulk RNA-seq, spatial transcriptomics, or pseudobulked single-cell compartments could potentially project each region or compartment independently onto the reference landscape, allowing within-tumor dispersion across the map to be measured directly.

Second, the integration of data from multiple independent studies and sequencing centers introduces potential technical variability. Although we applied uniform preprocessing and batch correction to mitigate study-specific effects and observed effective dataset mixing in the embedding, residual confounding cannot be completely excluded. Future work incorporating prospective, complementary single-cell or spatial profiling will further refine the resolution and robustness of the atlas. Finally, expression-based inference of potential therapeutic vulnerability does not establish functional dependency and requires experimental validation.

Third, the current landscape is RNA-defined and has not been validated as an immunohistochemistry-based pathology workflow. RNA-defined states may not directly correspond to protein-level staining patterns, and cluster identity should not be assigned in routine pathology practice using candidate IHC markers alone. Prospective validation with matched RNA-seq, tissue sections, standardized IHC protocols, scoring criteria, cutoffs, and centralized pathology review will be required to determine whether the lineage, neuroendocrine, proliferative, immune, and stromal programs identified here have reproducible protein-level correlates.

The survival analysis should be interpreted as hypothesis-generating rather than as validation of a clinically deployable prognostic classifier. Although A4 and SC were independently associated with survival after adjustment for cohort, sex, and smoking status, the model showed modest discrimination and was not benchmarked against a fully specified clinical model incorporating uniformly available stage, performance status, treatment received, treatment era, and oncogenotype. These variables were not consistently available across the publicly available datasets included in the integrated analysis. Future validation in clinically harmonized cohorts will be required to determine whether transcriptomic cluster assignment provides incremental prognostic information beyond established clinical predictors.

## STAR⍰METHODS

### KEY RESOURCES TABLE

| REAGENT OR RESOURCE | SOURCE | IDENTIFIER |
| --- | --- | --- |
| **Software and algorithms** |  |  |
| SRA-Toolkit 2.11.0 (fastq-dump) |  | https://trace.ncbi.nlm.nih.gov/Traces/sra/sra.cgi?view=software |
| R | R Foundation | https://www.r-project.org/ |
| R package “GSVA” (v 1.52.3) | Hanzelmann et al<sup>30</sup> |  |
| STAR2 (v2.7.7) | Dobin et al<sup>31</sup> |  |
| HTSeq (v0.11.0) | Anders et al<sup>32</sup> |  |
| ComBat-seq (sva package, v 3.52.0) | Zhang et al<sup>33</sup> |  |
| Arriba (v2.1.0) | Uhrig et al<sup>34</sup> |  |
| CaSpER | Harmanci et al.<sup>35</sup> |  |
| DESeq2 | Anders et al.<sup>36</sup> |  |
| EnrichR | Chen et al<sup>37</sup> |  |
| R package “survival” (v3.5.7) |  | https://github.com/therneau/survival |
| R package ConsensusClusterPlus ( v 1.68.0) | Wilkerson et al<sup>38</sup> |  |
| R package “ggplot2” ( v3.5.1) |  | https://cran.r-project.org/web/packages/ggplot2/ |
| Custom code used in this study |  | https://github.com/sonali-bioc/LungLandscapes |

### Data Collection and Processing

Publicly available bulk RNA sequencing datasets were obtained from GEO and GDC repositories. TCGA-LUAD, TCGA-LUSC, APOLLO-LUAD, and CPTAC-3 lung cancer datasets were obtained through the GDC Data Portal. Additional independent lung cancer cohorts were obtained from GEO. Datasets generated from cell lines, patient-derived xenografts, organoids, or other experimental model systems were excluded to maintain a patient-derived tumor transcriptomic framework. Studies using assay-specific or modified RNA-sequencing approaches that were not suitable for integration within the common bulk RNA-seq processing framework were also excluded.

Raw FASTQ files were uniformly processed to ensure cross-cohort consistency. Quality control was performed using FastQC (v0.11.9) and MultiQC (v1.9). Reads were aligned to the GENCODE GRCh38.primary_assembly reference genome using STAR (v2.7.7, two-pass mode). Gene-level counts were generated with HTSeq (v0.11.0) using GENCODE v39 annotations. Gene-level count matrices were aggregated across cohorts. Batch effects were corrected using ComBat-seq (sva), and variance-stabilized transformation (VST) values for only protein coding genes, were computed using DESeq2 for downstream visualization and analysis.

### Dimensionality Reduction and Clustering

Consensus clustering was performed independently in expression space using all protein coding genes and did not use two-dimensional embedding coordinates. Consensus clustering on full cohort formed three clusters, Adenocarcinoma samples falling in the second cluster were clustered separately using R package ConsensusClusterPlus (v 1.68.0) to get 5 clusters – A2, A3, A4, SC, and MD1. Squamous cell carcinoma samples were clustered in a separate analysis using consensus clustering to get two clusters – SQ1 and SQ2. Dimensionality reduction was performed using PCA, t-SNE, UMAP, and PaCMAP. PaCMAP embeddings was retained as the primary two-dimensional visualization because it provided a clear display of local sample neighborhoods and broader cross-histology relationships.

Landscape stability was assessed by subsampling 25%, 50%, 70%, and 90% of the full cohort and independently reconstructing PaCMAP embeddings using the same expression feature space and analysis parameters. Each embedding was annotated by dataset, histological diagnosis, and transcriptional cohort to assess preservation of the reference landscape structure across sampling fractions.

### Clinical, molecular, and tumor microenvironment annotation of transcriptional clusters

Sample-level clinical, molecular, and tumor microenvironment annotations were assembled for tumors included in the lung cancer transcriptional landscape and linked to transcriptional cluster assignment. Clinical annotations included pathologist-assigned histology, smoking status, sex, AJCC pathologic stage, and specimen type. Histology was harmonized as lung adenocarcinoma (LUAD), lung squamous cell carcinoma (LUSC), small cell lung cancer (SCLC), or non-small cell lung cancer not otherwise specified (NSCLC-NOS). Smoking status was harmonized where available; pack-year information was not uniformly available and was therefore not included in quantitative smoking-exposure analyses.

DNA-based somatic mutation calls were used to annotate clinically relevant driver genes in samples with matched DNA data from TCGA-LUAD, TCGA-LUSC, and CPTAC-3. Samples were classified as mutation-positive for a gene if at least one somatic mutation was detected in that gene and mutation-negative otherwise; samples without matched DNA data were labeled as no data. Driver genes included EGFR, KRAS, ALK, ROS1, RET, MET, BRAF, and ERBB2/HER2.

Nonsynonymous mutation burden was calculated among samples with matched DNA mutation data after restricting variants to nonsynonymous consequences, including missense, stop-gained, stop-lost, start-lost, frameshift, in-frame insertion or deletion, and splice acceptor or donor variants. Because callable territory was not uniformly available across cohorts, mutation burden was reported as the number of nonsynonymous somatic mutations per sample rather than as mutations per megabase.

For cluster-level oncogenotype annotation, we generated a combined oncogene status for each clinically relevant driver gene. For EGFR, KRAS, and BRAF, combined status was based on DNA mutation status. For ALK, ROS1, RET, MET, and ERBB2/HER2, combined status integrated DNA mutation and RNA fusion evidence. A sample was classified as combined oncogene-positive if either a DNA mutation or RNA fusion was detected for that gene, combined oncogene-negative if the relevant available assays were negative, and no data only when no relevant assay information was available. DNA mutation calls and RNA fusion calls were retained as separate evidence layers in addition to the combined oncogene status.

Tumor microenvironment annotations were derived using multiple approaches. Tumor purity was represented using PUREE purity estimates. Immune and stromal content were represented using ESTIMATE immune score, ESTIMATE stromal score, ESTIMATE score, xCell immune score, xCell stromal score, and xCell microenvironment score. CIBERSORTx immune cell fractions were included as cell-type-level immune deconvolution features, and PD-L1 expression was represented by log2-transformed CD274 TPM expression. Neuroendocrine marker status was summarized using ASCL1, CHGA, SYP, NCAM1, and INSM1 expression; each marker was standardized across samples, and a composite neuroendocrine marker score was calculated as the mean standardized expression across available markers.

Cluster-level associations were evaluated for each clinical and molecular variable. Categorical variables were summarized as cluster-by-variable contingency tables and tested using chi-square tests when expected cell counts were adequate or Fisher’s exact tests with simulated p-values for sparse tables. Cluster-specific enrichment was assessed using cluster-versus-rest Fisher’s exact tests. Continuous variables, including nonsynonymous mutation burden, neuroendocrine marker score, tumor purity, immune and stromal scores, CIBERSORTx fractions, xCell scores, ESTIMATE scores, and CD274 expression, were compared across clusters using Kruskal-Wallis tests followed by cluster-versus-rest Wilcoxon rank-sum tests. P-values were adjusted using the Benjamini-Hochberg false discovery rate procedure. For each analysis, denominators were calculated using samples with available data for the corresponding variable, and unavailable clinical or assay-specific data were retained explicitly rather than coded as negative calls.

### Comparison with established molecular subtype classifications

Established molecular subtype annotations were compared with the nine transcriptional landscape clusters for LUAD, LUSC, and SCLC. For LUAD and LUSC, published subtype labels were retained when available from the source cohorts. In parallel, subtype assignments were inferred across the integrated cohort using published subtype-associated gene signatures: bronchoid/magnoid/squamoid for LUAD and basal/classical/primitive/secretory for LUSC. For SCLC, samples were classified according to the SCLC-A, SCLC-N, SCLC-P, and SCLC-I framework using correlation to published subtype expression profiles, with low-confidence assignments retained as ambiguous. Subtype distributions were summarized within each landscape cluster as stacked percentages across all samples in that cluster, preserving the cross-histology composition of the integrated landscape. Samples without an applicable inferred disease-specific subtype assignment were labeled “Not Classified,” whereas samples lacking a published subtype annotation were labeled “Not Reported.”

Subtype-cluster concordance was quantified separately for LUAD, LUSC, and SCLC using disease-matched samples with available reported or inferred subtype assignments. Cluster-by-subtype contingency tables were constructed for each subtype framework. Overall association between landscape cluster and subtype assignment was evaluated using contingency-table testing, with Cramer’s V used as an effect-size measure. Agreement between landscape clusters and prior subtype systems was additionally summarized using adjusted Rand index and normalized mutual information. Enrichment of individual subtype programs within specific landscape clusters was evaluated using Fisher’s exact tests followed by Benjamini-Hochberg correction.

### Differential Expression and Pathway Analysis

Differential gene expression analysis was performed using DESeq2, with significance defined as FDR < 0.05 and |log fold change| > 0.5. For cluster level analysis, genes upregulated in clusters A2–A4,SC, MD1, and MD2 were identified relative to A1, which served as the reference group. Genes significantly downregulated relative to A1 were aggregated to define transcriptional programs enriched within the A1 cluster. All differentially expressed genes from each cluster were analyzed using EnrichR to determine the biological signature of the cluster.

In an independent analysis, Gene sets from KEGG, Biocarta, Reactome, and GO Biological Processes (MSigDB v7.2) were analyzed using GSVA on batch-corrected VST values, generating pathway activity scores (−1 to 1). Pathway scores were visualized over PaCMAP using ggplot2.

### Fusion and Copy Number Inference

Gene fusions were detected using Arriba (v2.1.0) and STAR-Fusion on STAR two-pass alignments. Only high-confidence fusions identified by both tools and involving at least one protein-coding gene (GENCODE v39, GRCh38.p14) were retained.

Large-scale copy number alterations were inferred from bulk RNA-Seq using CaSpER. BAFExtract was used for B-allele frequency pileups, and hg38 cytoband/centromere annotations were obtained from UCSC.

### Survival Analysis

Kaplan–Meier analyses were performed using samples with available overall survival time and event data. Significance was assessed using the survival R package (v3.5.7). Multivariable Cox models were adjusted for relevant covariates as specified in Results. To evaluate the robustness of survival associations independent of reference selection, pairwise comparisons among all clusters were subsequently performed using estimated marginal means with Tukey adjustment for multiple testing.

### Projection of new samples onto the reference landscape

To project a new tumor sample onto the lung cancer transcriptomic landscape, raw bulk RNA-seq FASTQ files should be processed using the same reference workflow used to construct the atlas. Reads are aligned to the GENCODE GRCh38 primary assembly reference genome using STAR2 (v2.7.7a), and gene-level counts are quantified using HTSeq (v0.11.0) with GENCODE v39 gene annotations. Counts are restricted to GENCODE v39 protein-coding genes and transformed using variance-stabilizing transformation (VST) normalization in DESeq2. The VST-normalized expression matrix from the reference landscape is then downloaded from the Lung Landscapes resource, and the incoming sample is aligned to the same protein-coding gene feature space and gene order.

For the public workflow, the incoming sample is aligned to the reference protein-coding gene feature space and projected into the existing PaCMAP coordinate system using transform. The PaCMAP transform workflow is provided in the GitHub repository and does not require use of the patent-pending nearest-neighbor approach.

A laboratory-developed nearest-neighbor landing approach was also used for comparison in Figure S7B, in which incoming samples were compared with reference tumors in high-dimensional VST-normalized expression space and positioned relative to transcriptionally similar reference samples; this approach is not required for the public PaCMAP transform-based workflow.

### Oncoscape Integration

Expression and clinical matrices were formatted for cBioPortal compatibility and uploaded into Oncoscape with predefined visualization states corresponding to manuscript figures.

## Supplementary Table Legends

Table S1. Number of samples contributed by each dataset included in the integrated lung cancer transcriptomic landscape.

Table S2. Tukey-adjusted pairwise Cox proportional hazards comparisons across the nine landscape clusters, including hazard ratios, 95% confidence intervals, and adjusted P values.

Table S3. Chromosome arm–level copy number gains and losses stratified by lung cancer disease type.

Table S4. Distribution of clinically relevant fusion events across histologic disease categories and landscape clusters, including disease-and cluster-level fusion counts and percentages.

Table S5. Cluster-level association results for histology, sex, smoking status, stage, specimen type, mutation burden, mutations, fusions, neuroendocrine marker score, PUREE purity, ESTIMATE, xCell, CIBERSORTx, and CD274 expression.

Table S6. Quantitative comparison of landscape clusters with reported or inferred LUAD, LUSC, and SCLC subtype assignments, including global association tests, Cramer’s V, ARI, NMI, and cluster-specific enrichment statistics.

Table S7. Differential gene expression analysis results across transcriptional clusters.

Table S8. Pathway enrichment results for adenocarcinoma clusters A1–A4 and the mixed-diagnosis region MD1.

Table S9. Pathway enrichment results comparing the small cell lung cancer cluster SC and SCLC diagnosed tumors localized within MD2.

Table S10. Pathway enrichment results for SQ1 and SQ2 squamous carcinoma clusters.

Table S11. Coordinates and metadata for patient tumors and patient-derived xenograft (PDX) samples projected onto the reference PaCMAP landscape.

Table S12. RNA-nominated cluster-associated programs and suggested functional validation strategies, including perturbational, pharmacologic, model-system, and treatment-response studies.

## Acknowledgements

We thank Thomas J. Lynch and members of the Holland lab at Fred Hutch Cancer Center for valuable discussions and review of the manuscript. This research was supported by funding from the Fred Hutch Cancer Center (E.C.H), 1R35 CA253119-01A1 (E.C.H).

## Funding

National Institutes of Health grant 1R35 CA253119-01A1 (E.C.H).

## Author Contributions

Conceptualization, S.A., and E.C.H.; Methodology, S.A. and E.C.H.; Formal Analysis, L.S., N.H. and S.A.; Software, M.J., G.G; Investigation, S.A., J.F., D.M., A.H.B and E.C.H.; Resources, D.M. and A.B.; Data Curation, S.A., H.N. and L.S., Writing – Original Draft, S.A and E.C.H; Writing – Review & Editing, S.A., E.C.H, C.C.P, E.Q.K, A.H.B., D.M.; Visualization, S.A., L.S., M.J., G,G; Supervision, E.C.H., Funding Acquisition, E.C.H.

## Conflict of interest

A patent application related to a nearest-neighbor sample-landing approach has been filed and S.A, M.J and E.C.H are listed as inventors (Serial No.: 63/960,422). This application is pending and has not been issued as a patent. The patent-pending component does not cover the public datasets, atlas construction workflow, consensus clustering, dimensionality-reduction analyses, clinical and molecular annotation analyses, survival analyses, or the PaCMAP transform-based projection workflow described in this manuscript.

## Data and materials availability

All analysis including statistics and visualization were done in R version 4.4.0. Plots were generated using R basic graphics and ggplot2. Raw sequencing data was downloaded from GEO as shown in Table S1. All custom code used in this study is available at https://github.com/sonali-bioc/LungLandscapes. To preserve reproducibility, we provide a public analysis workflow and an open PaCMAP transform projection option; the patent-pending component is limited to the laboratory-developed nearest-neighbor landing approach and is not required to reproduce the analyses presented here.

## Ethics

This study was conducted using publicly available and fully de-identified transcriptomic datasets. For new data collected from human subjects, institutional review board (IRB) approval and informed consent was required. All original studies from which data were obtained had received appropriate ethical approvals and consent for data sharing.

## Supplemental Figures

**Supplementary Figure 1.**
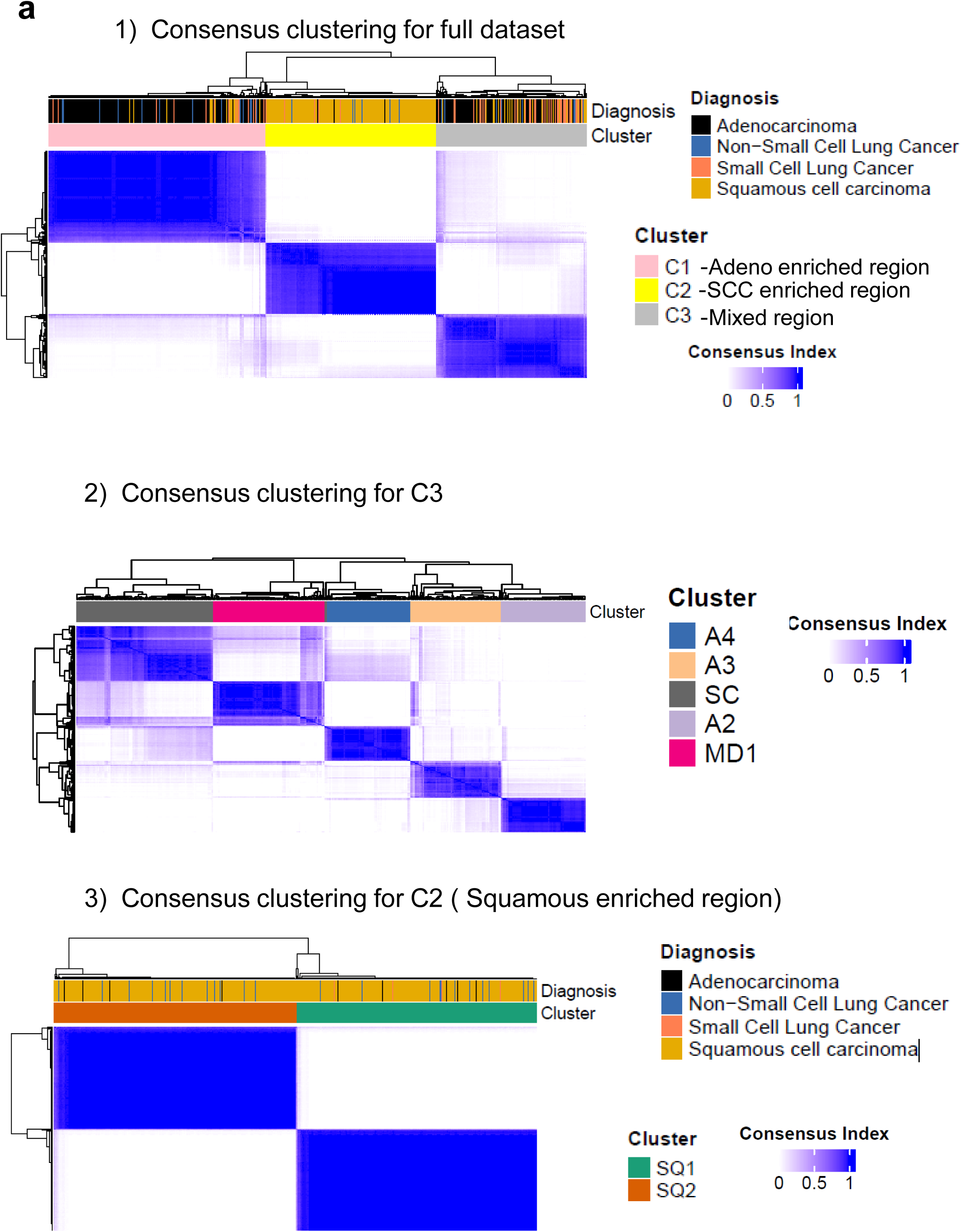
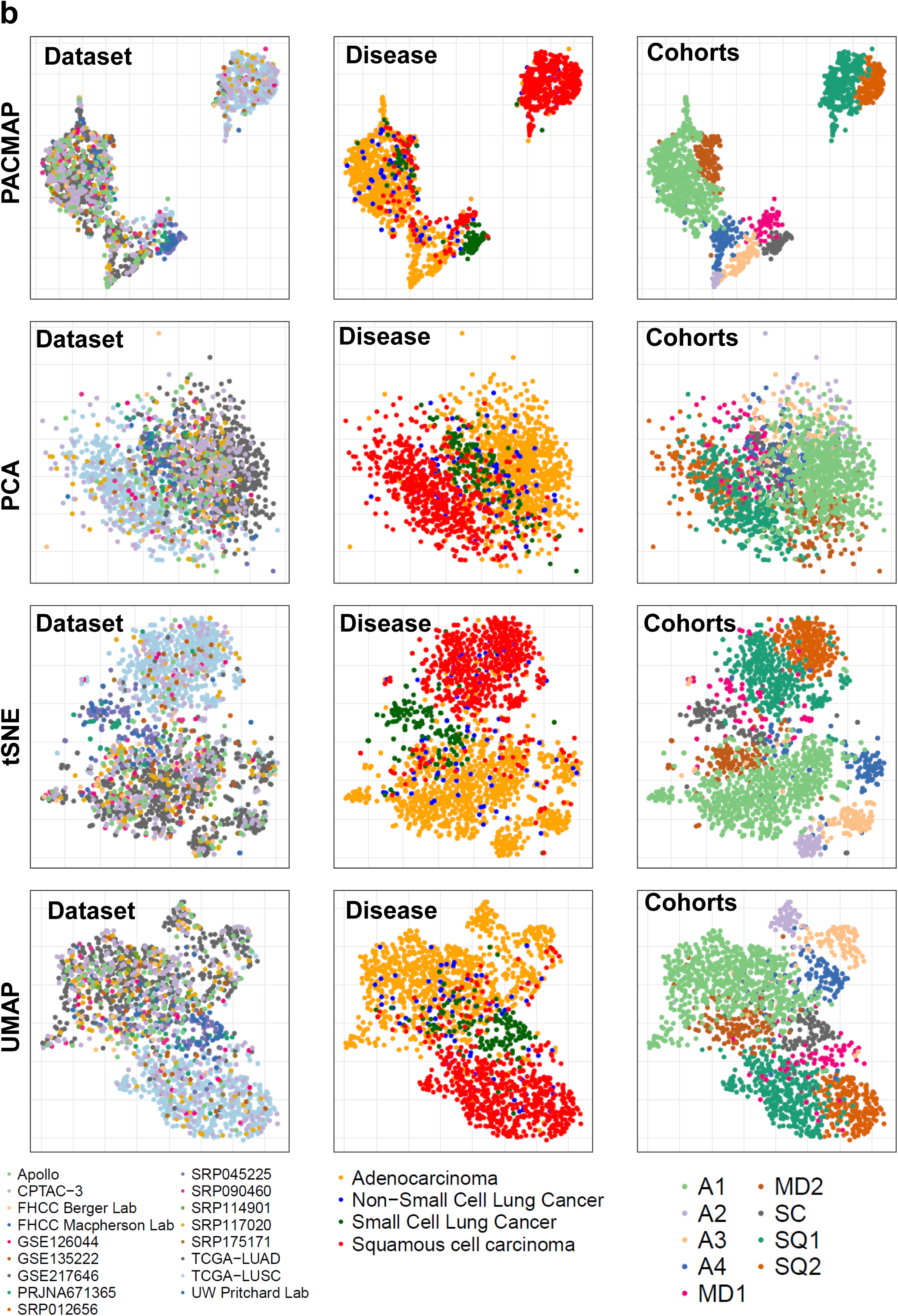
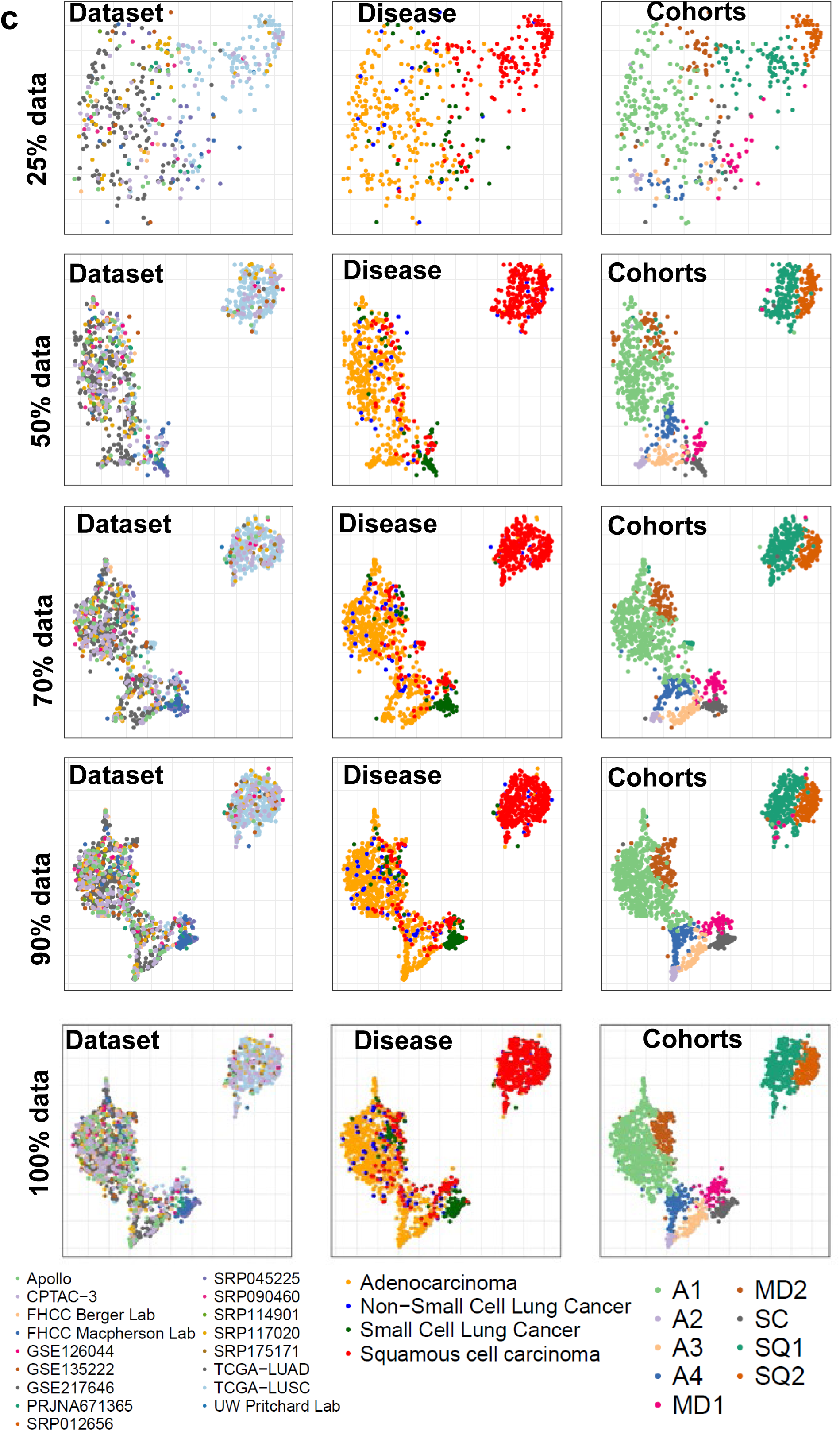
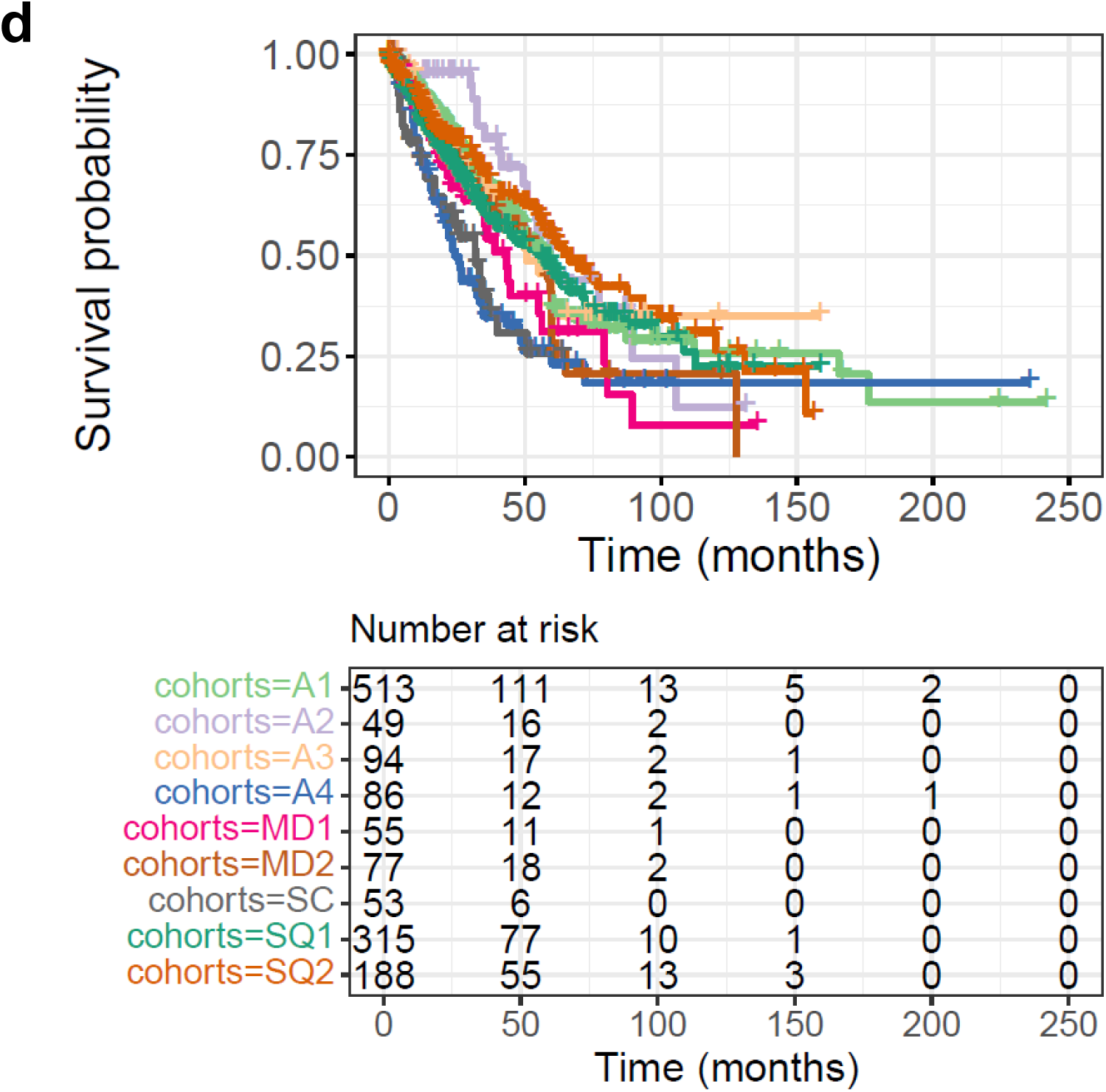
Additional annotations and clustering analyses of the lung cancer landscape. (A) Consensus clustering heatmap of (1) full cohort (2) samples restricted to the second mixed region. (3) samples restricted to pure squamous region (B) PCA, UMAP, t-SNE, and PaCMAP were generated using the same batch-corrected, variance-stabilized expression matrix comprising all protein-coding genes. For each method, samples are shown colored by disease annotation, dataset of origin, and consensus-cluster assignment. (C) PaCMAP embeddings were reconstructed after random subsampling of 25%, 50%, 70%, and 90% of the full cohort. For each subsampling level, samples are shown colored by disease annotation, dataset of origin, and consensus-cluster assignment. (D) Kaplan–Meier survival curves across cohorts with the corresponding risk table shown below the plot.

**Supplementary Figure 2.**
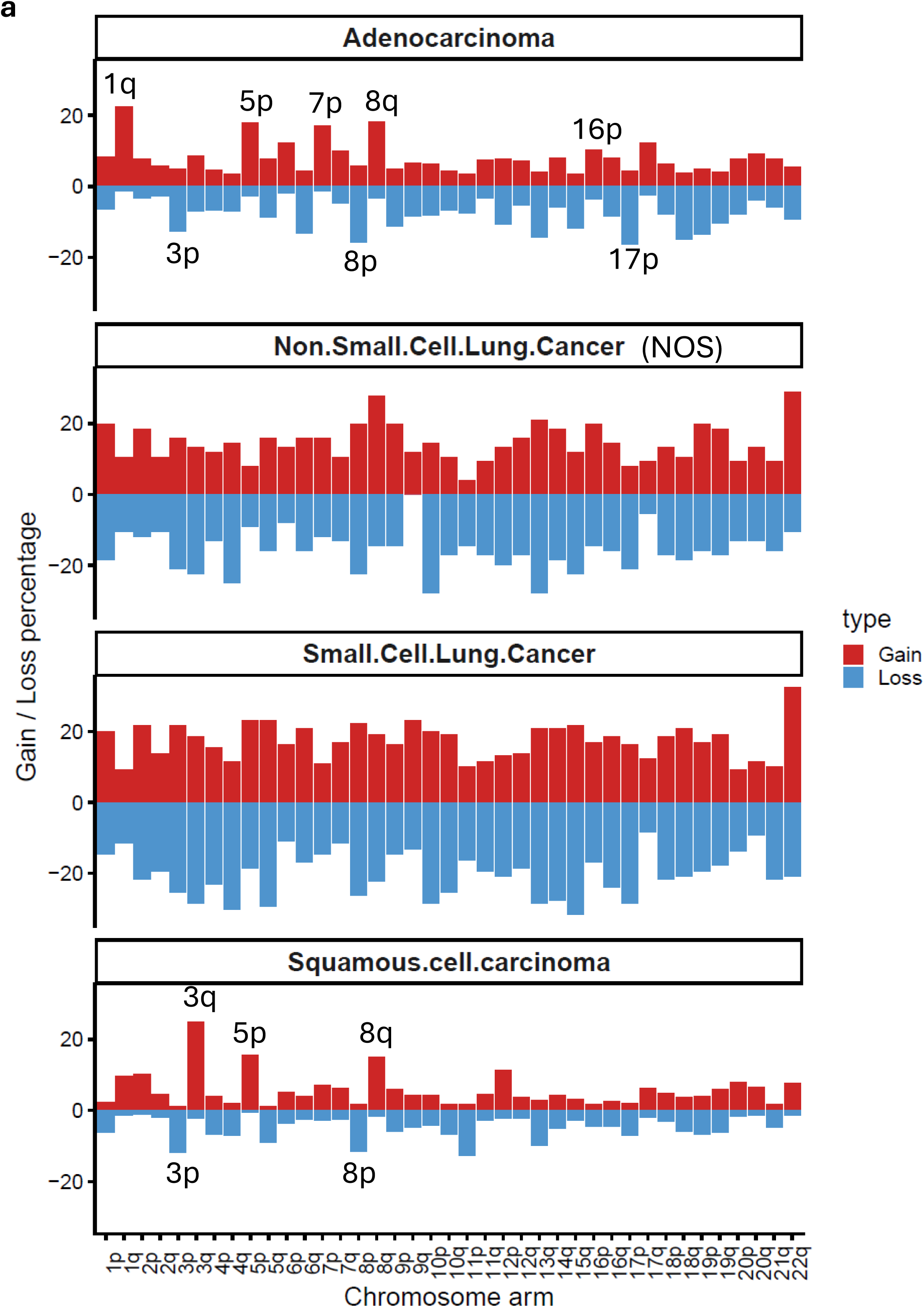
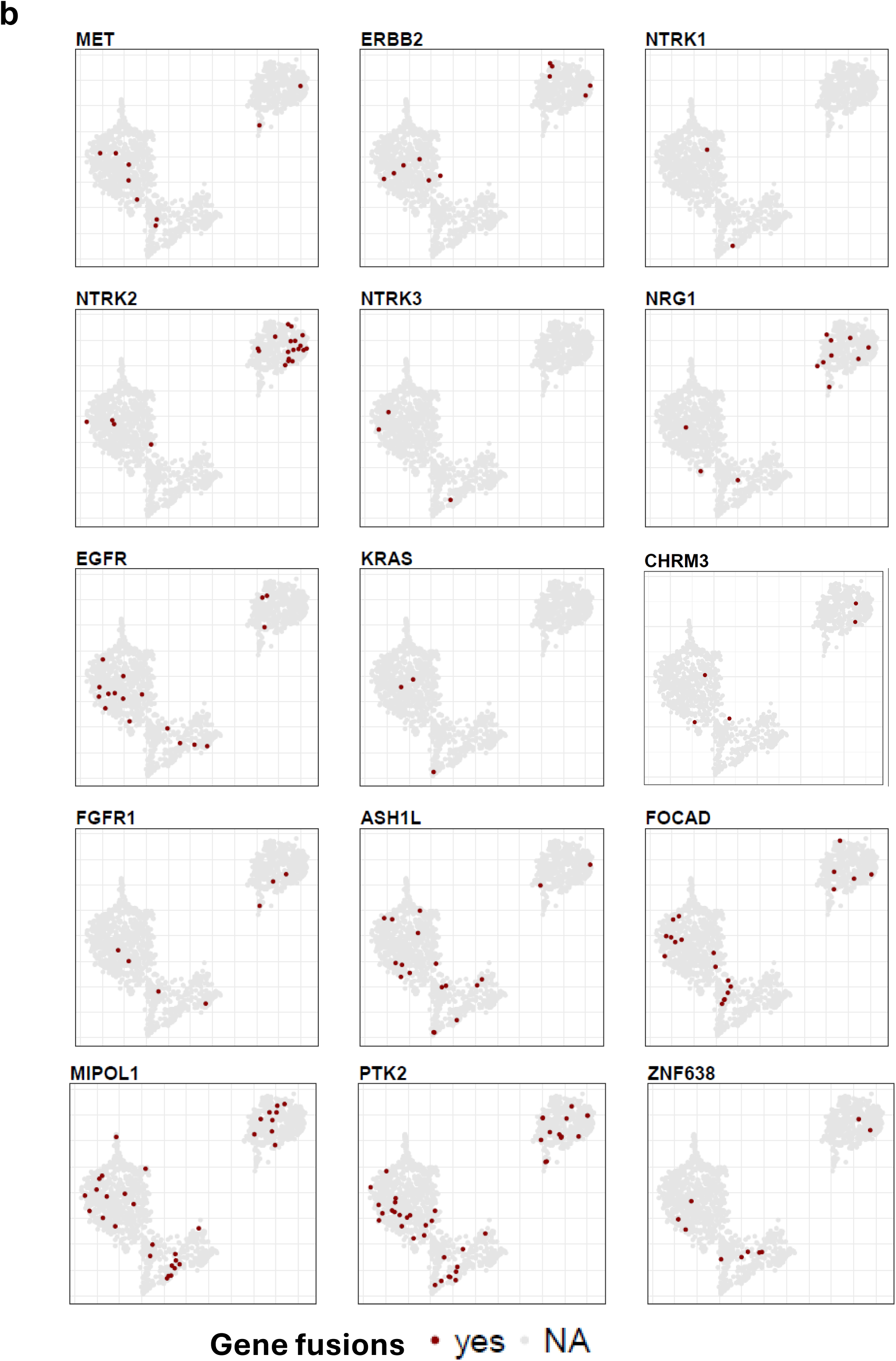
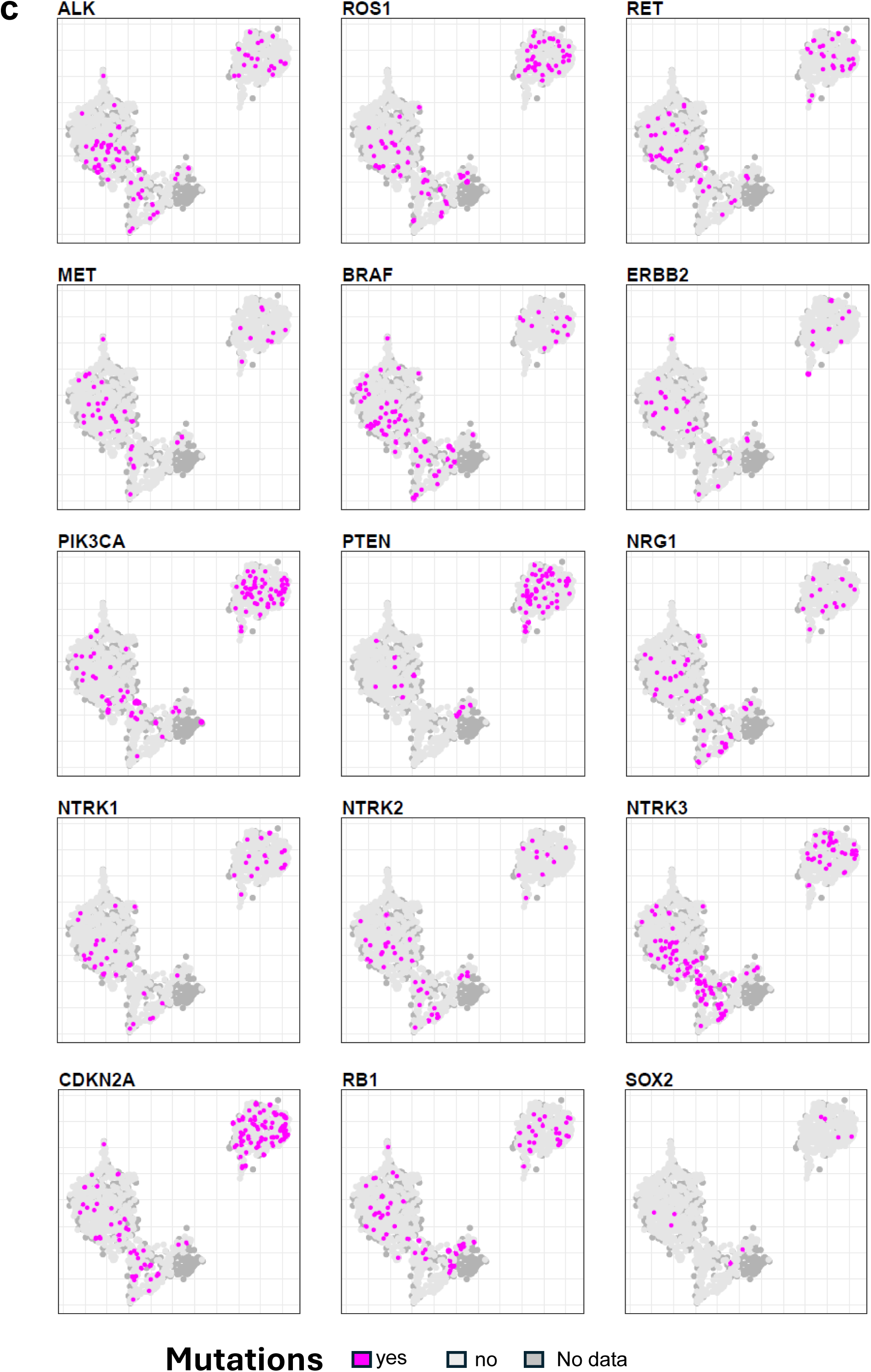
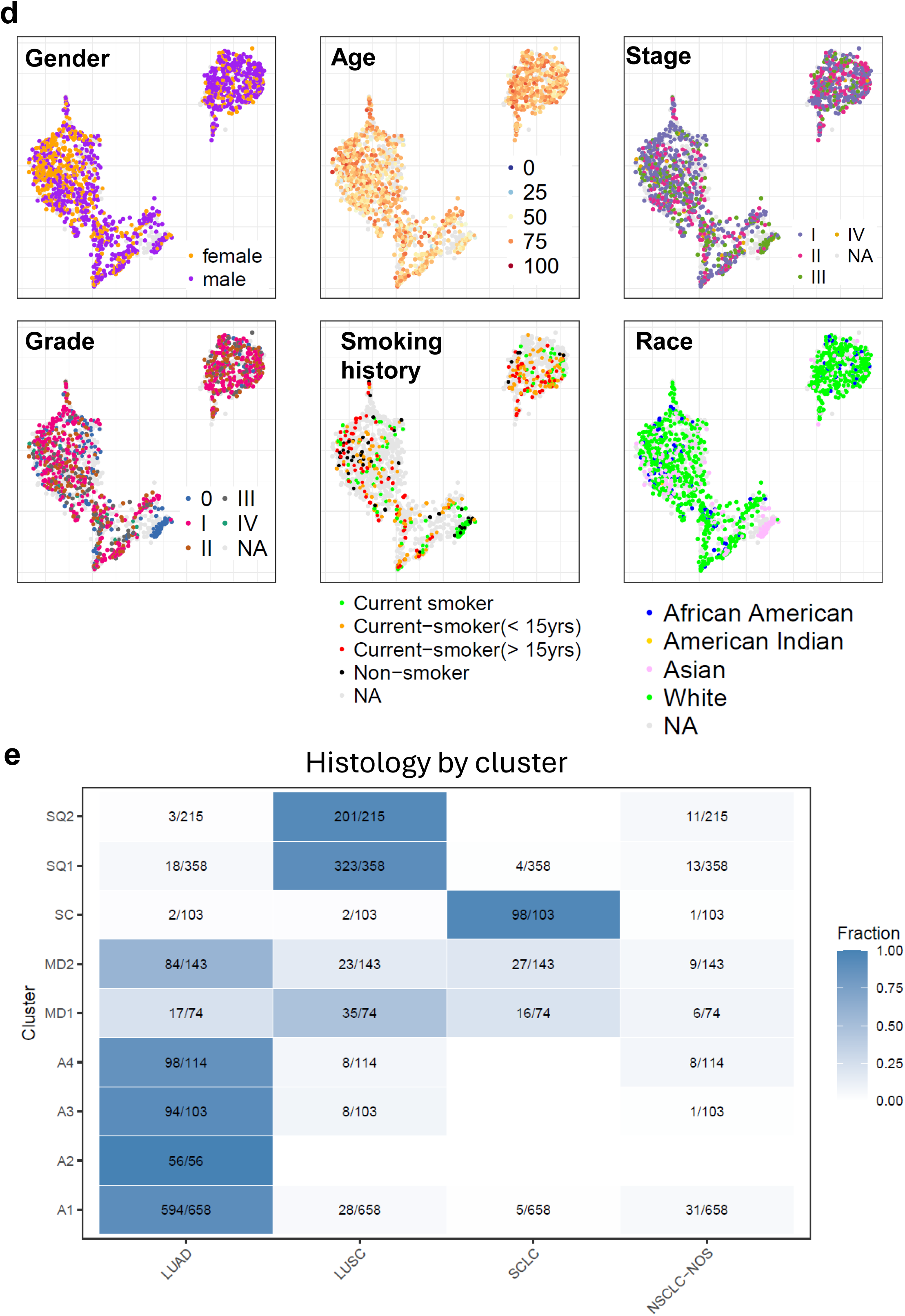
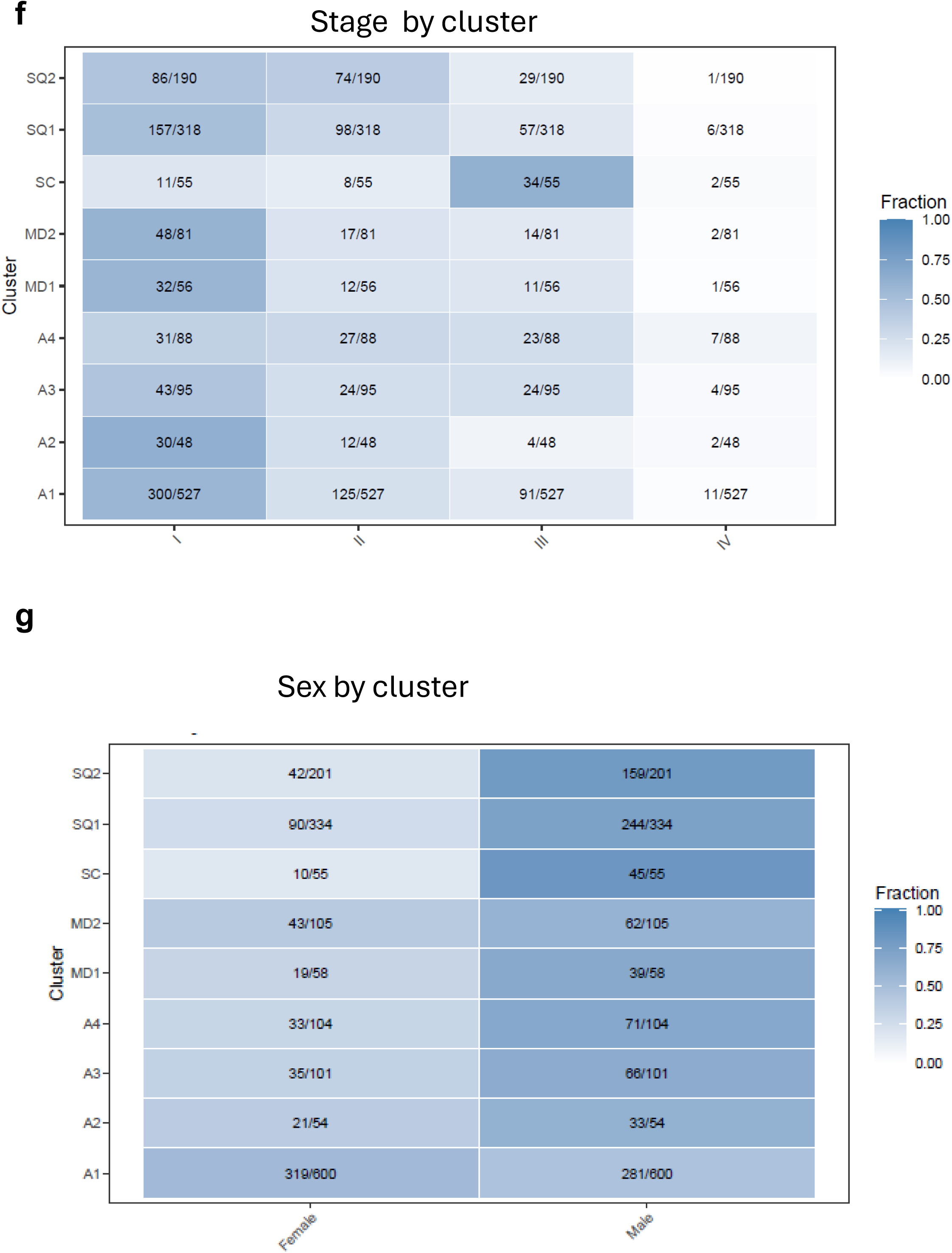
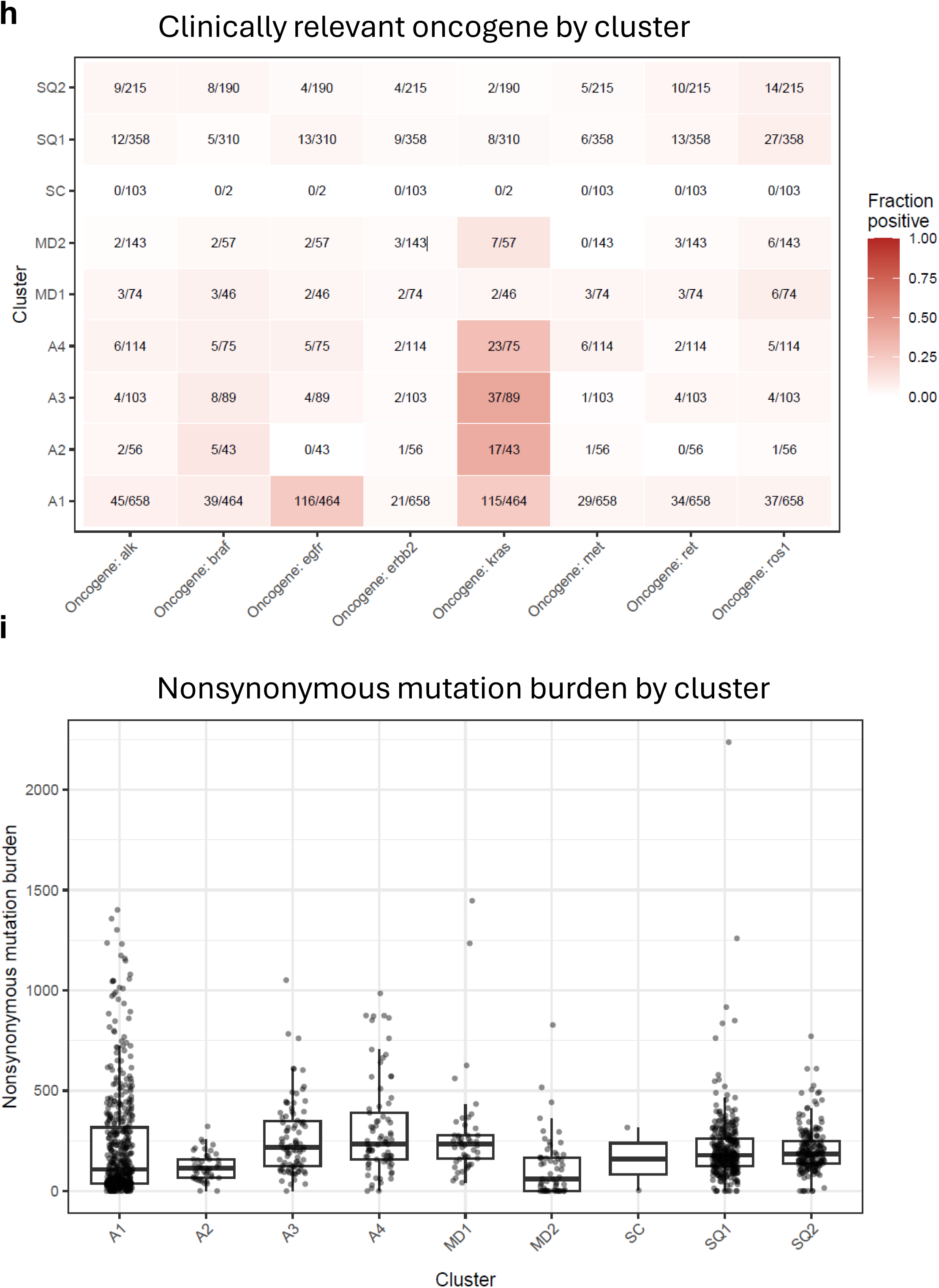
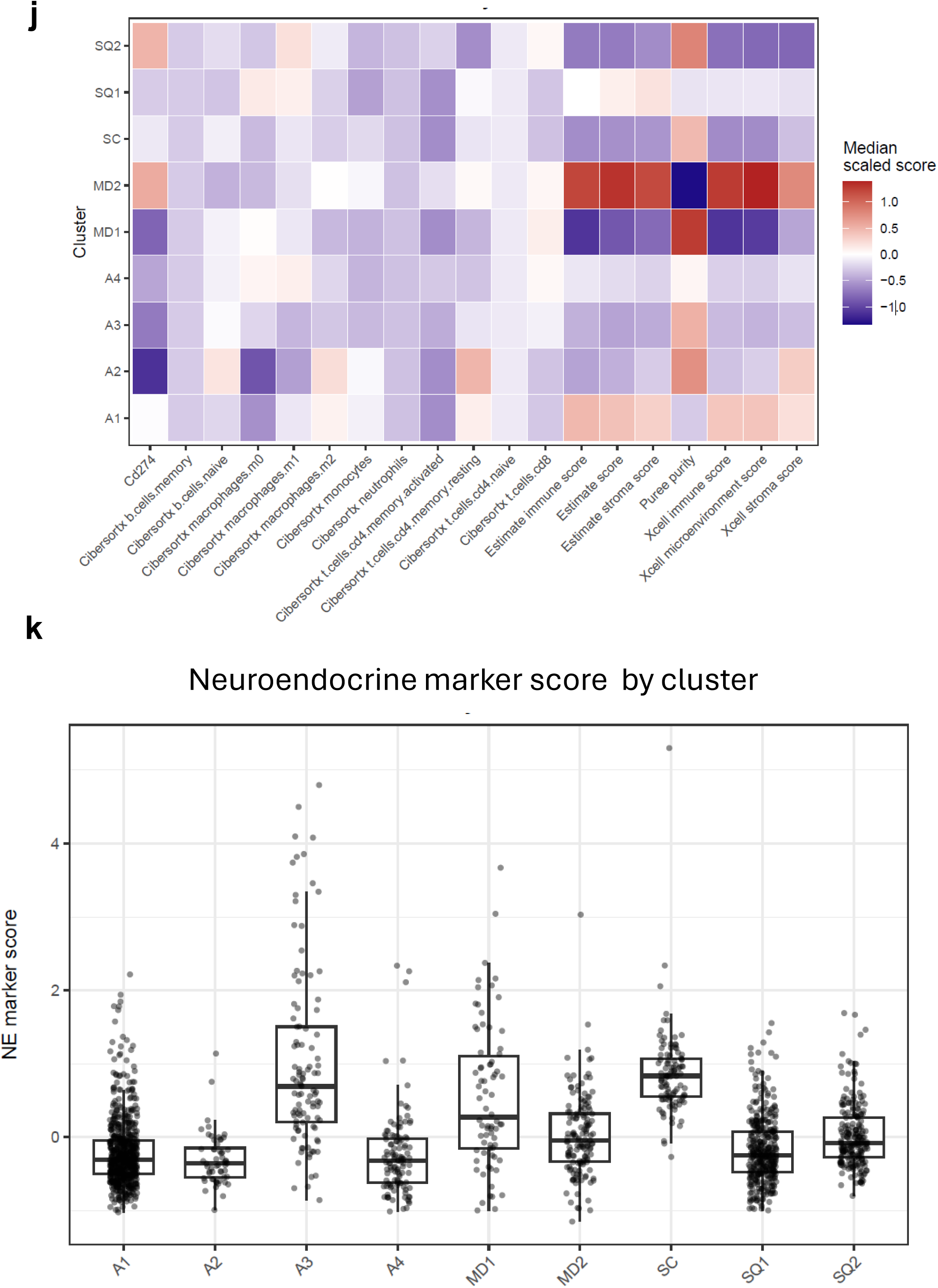
Copy number alterations and gene fusions across the lung cancer landscape. (A) Manhattan plot summarizing chromosome arm–level copy number alterations across diseases. (B) PaCMAP embedding highlighting samples harboring gene fusions involving MET, NTRK2, FGFR1, ASH1L, FOCAD, MIPOL1, PTK2, ZNF638, and CHRM3. Samples with detected fusions are colored by gene, while all other samples are shown in grey. (C) PaCMAP embedding highlighting samples harboring mutations. Samples with detected mutations are colored by gene, while all other samples are shown in grey. The samples with no-matched DNA calls are colored in darkgrey and were printed first. (D) PaCMAP embeddings colored by clinical variables: pathological stage, grade, age at diagnosis, race, sex, and smoking history. Figure S2E–K. Clinical, molecular, and tumor microenvironment annotation of transcriptional clusters. (E) Distribution of pathologist-assigned histology across transcriptional clusters. Histology was harmonized as LUAD, LUSC/SCC, SCLC, or NSCLC-NOS. (F) AJCC pathologic stage distribution by transcriptional cluster among tumors with available stage annotation. (G) Sex distribution by transcriptional cluster among tumors with available sex annotation. (H) Cluster-level distribution of clinically relevant oncogene alterations, integrating DNA mutation and RNA-derived fusion evidence where applicable. (I) Nonsynonymous mutation burden by transcriptional cluster among tumors with matched DNA mutation data. Mutation burden is reported as nonsynonymous mutation count rather than mutations per megabase because callable territory was not uniform across cohorts. (J) Heatmap summarizing tumor microenvironment features by cluster, including CD274 expression, CIBERSORTx immune cell fractions, xCell immune and stromal scores, ESTIMATE immune and stromal scores, and PUREE tumor purity. (K) Composite neuroendocrine marker score by transcriptional cluster, calculated from standardized expression of ASCL1, CHGA, SYP, NCAM1, and INSM1.

**Supplementary Figure 3.**
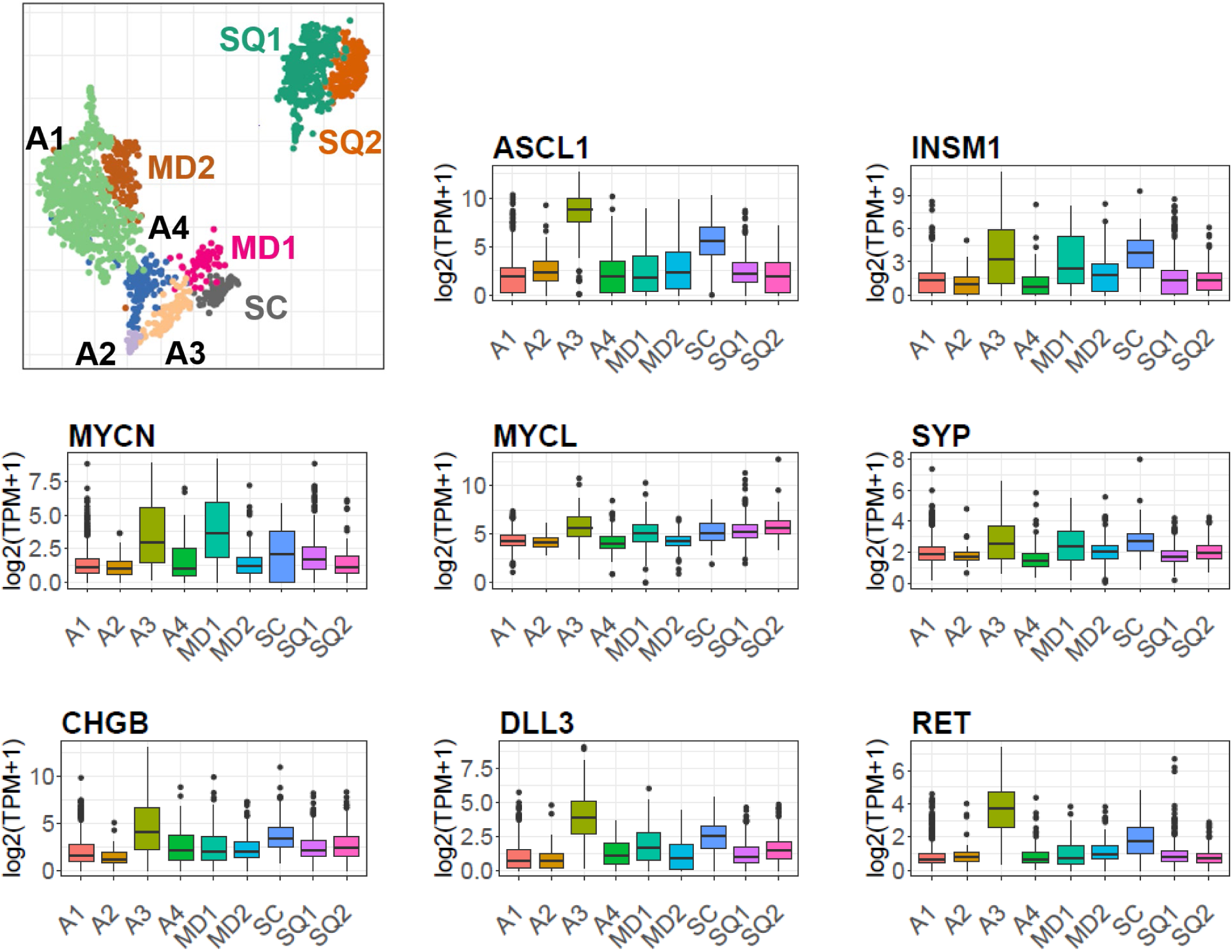
Gene expression patterns across transcriptional clusters. Schematic representation of the transcriptional clusters identified in the unified lung cancer landscape. Boxplots showing expression of representative genes across clusters, illustrating cluster-specific transcriptional signatures.

**Supplementary Figure 4.**
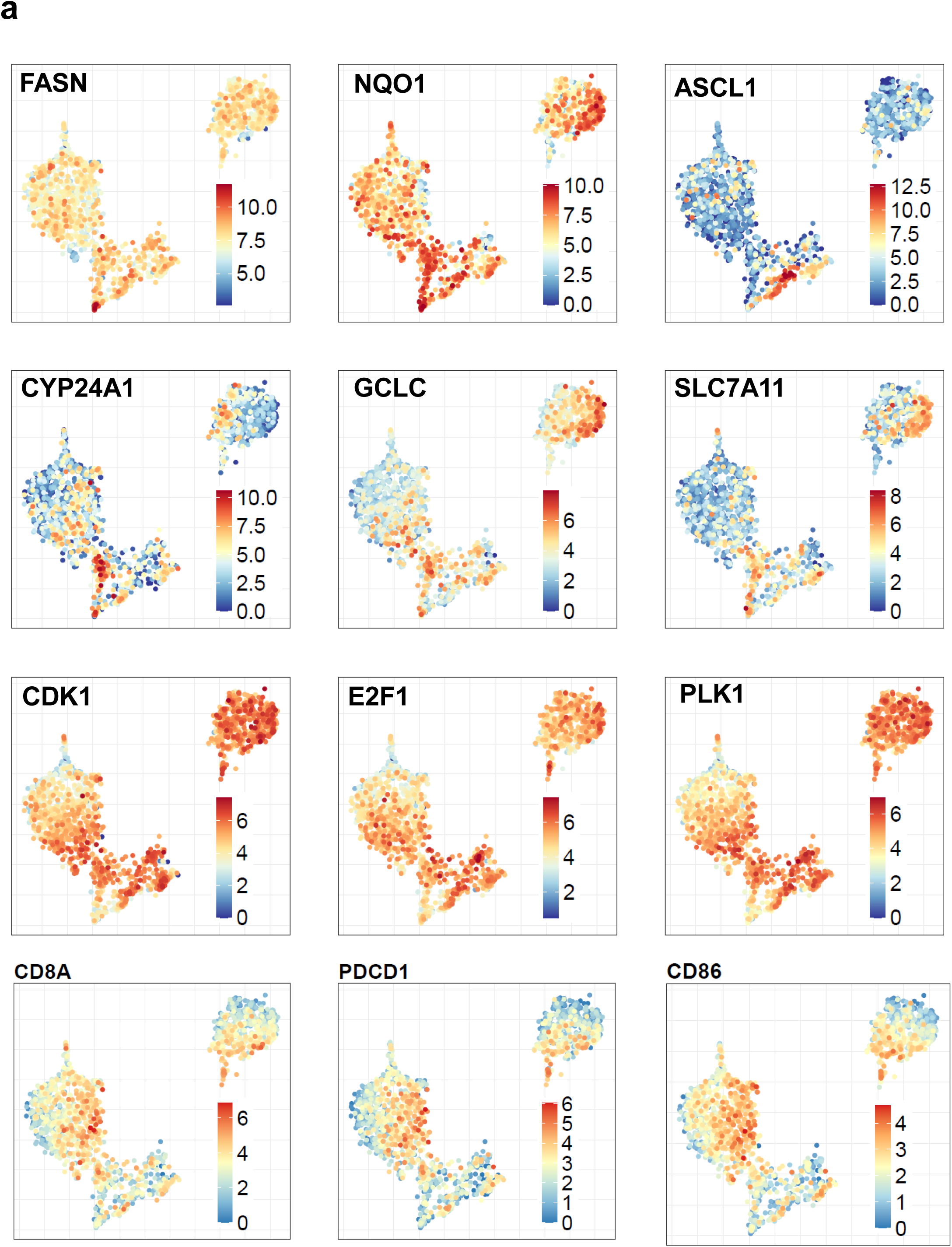
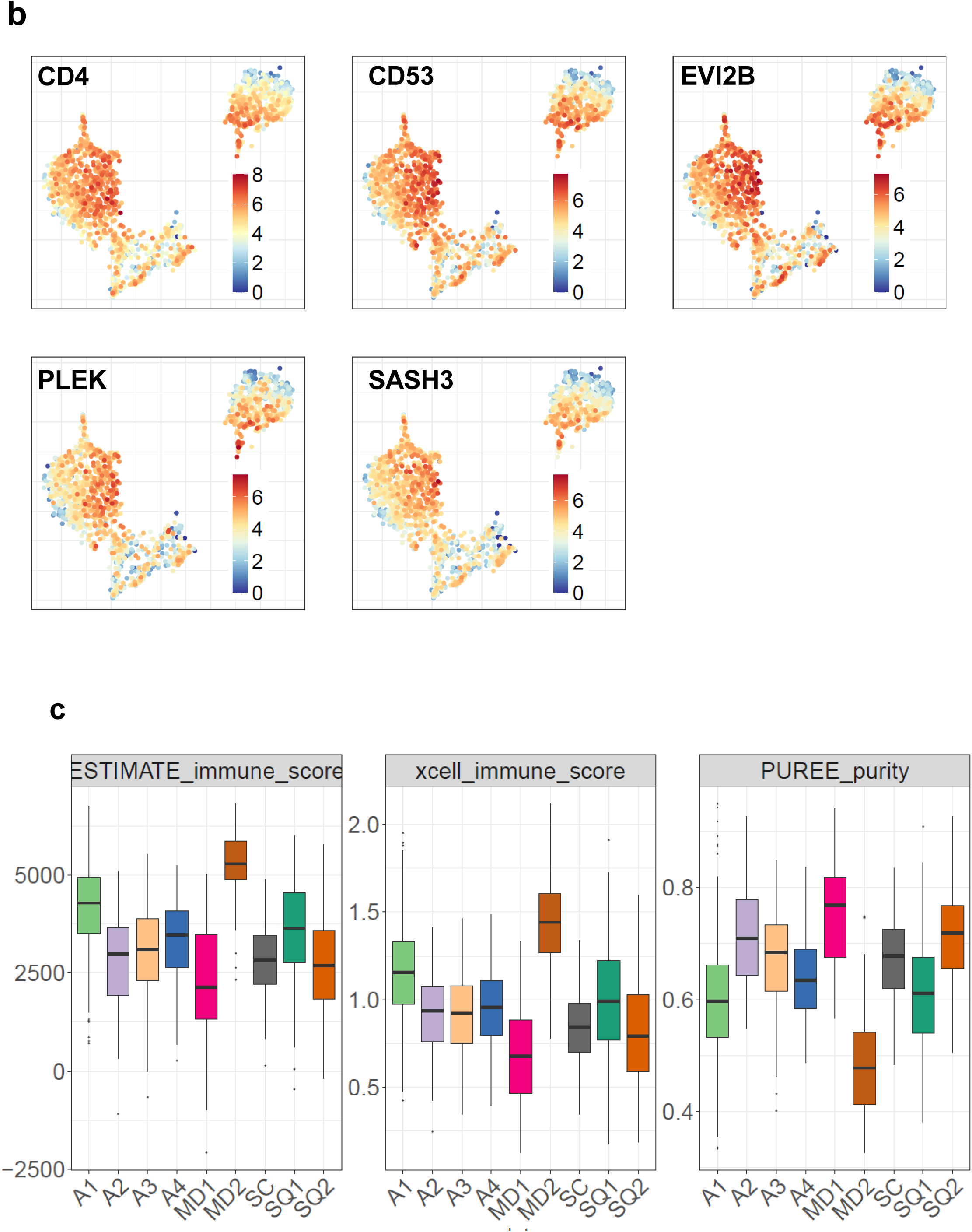
Immune-associated gene expression and tumor purity across clusters. (A) PaCMAP embeddings colored by expression of genes upregulated in the different transcriptomic clusters (B) PaCMAP embeddings colored by expression of genes comprising the five-gene immune predictor signature (CD4, CD53, EVI2B, PLEK, and SASH3), all enriched within the immune-infiltrated A1 and MD2 regions. (C) Boxplots showing immune score from ESTIMATE and Xcell, and tumor purity estimates from PUREE across clusters.

**Supplementary Figure 5.**
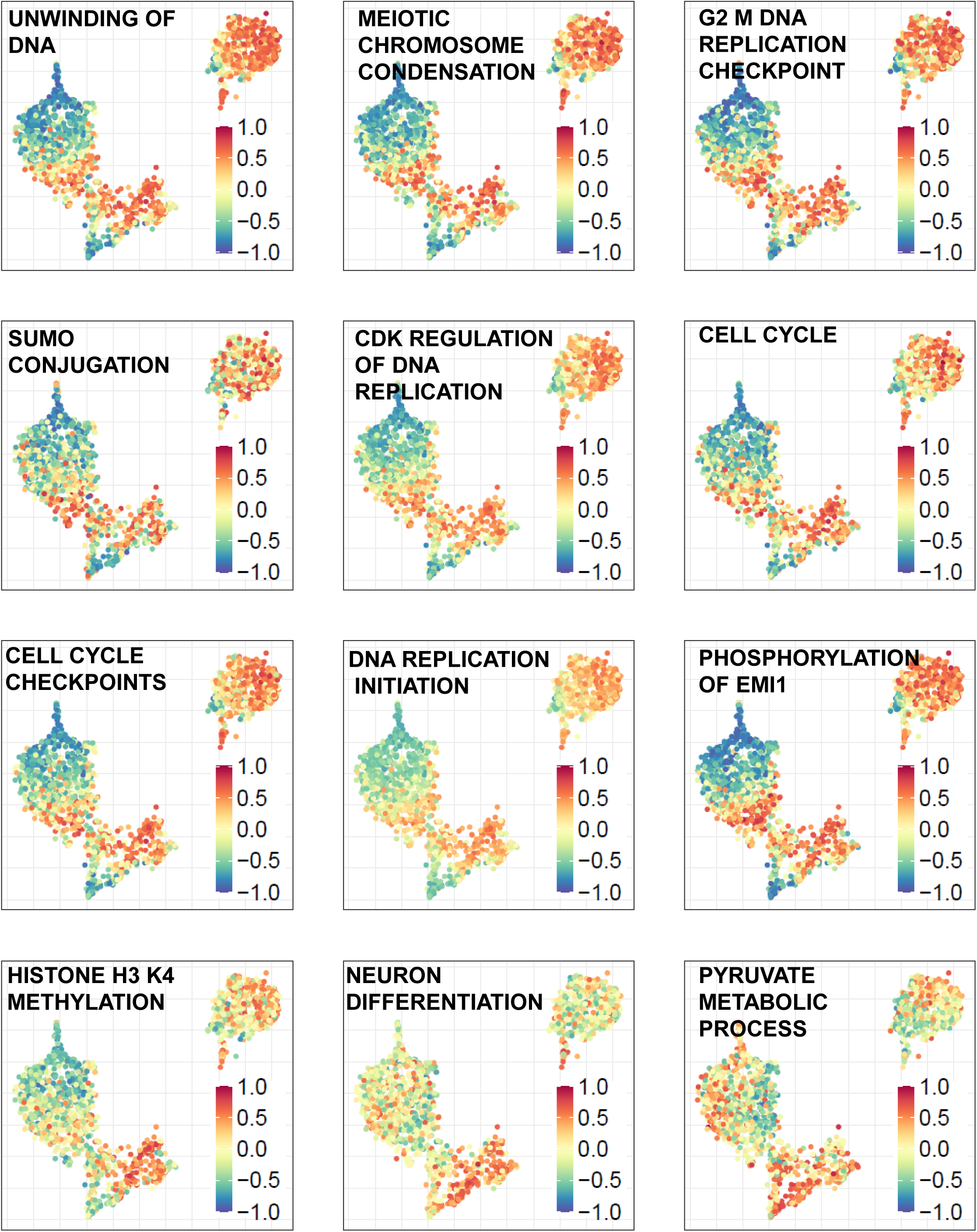
Pathway enrichment distinguishing small cell lung cancer states. PaCMAP embeddings colored by GSVA pathway scores for pathways enriched in the SC small cell lung cancer cluster and in SCLC-diagnosed tumors localized within MD2, illustrating distinct transcriptional programs between these regions.

**Supplementary Figure 6.**
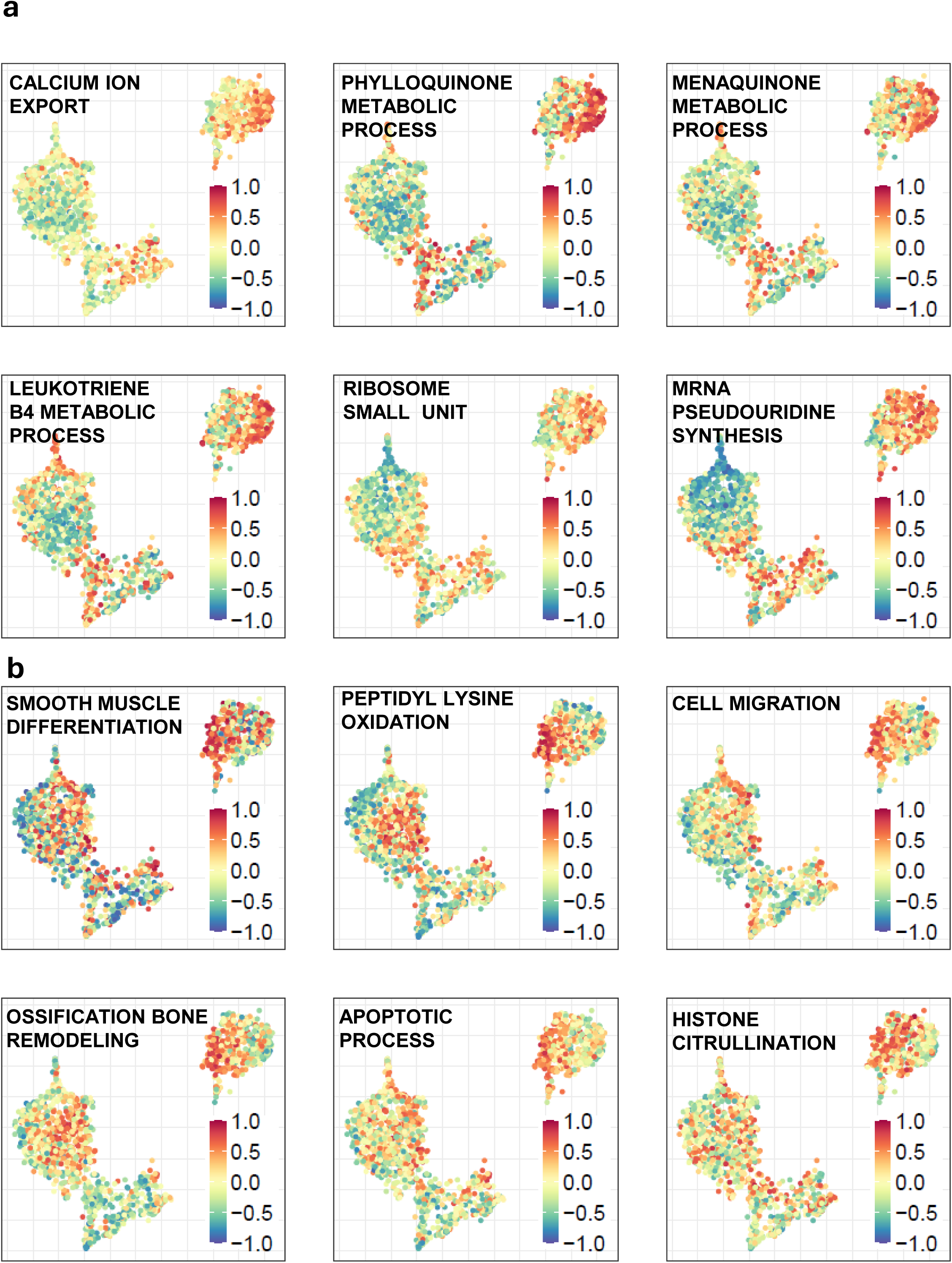
Pathway enrichment distinguishing squamous carcinoma states. PaCMAP embeddings colored by GSVA pathway scores for pathways enriched in SQ1 and SQ2, illustrating distinct transcriptional programs separating tumor-intrinsic metabolic/redox-adapted and immune-infiltrated squamous carcinoma states

**Supplementary Figure 7.**
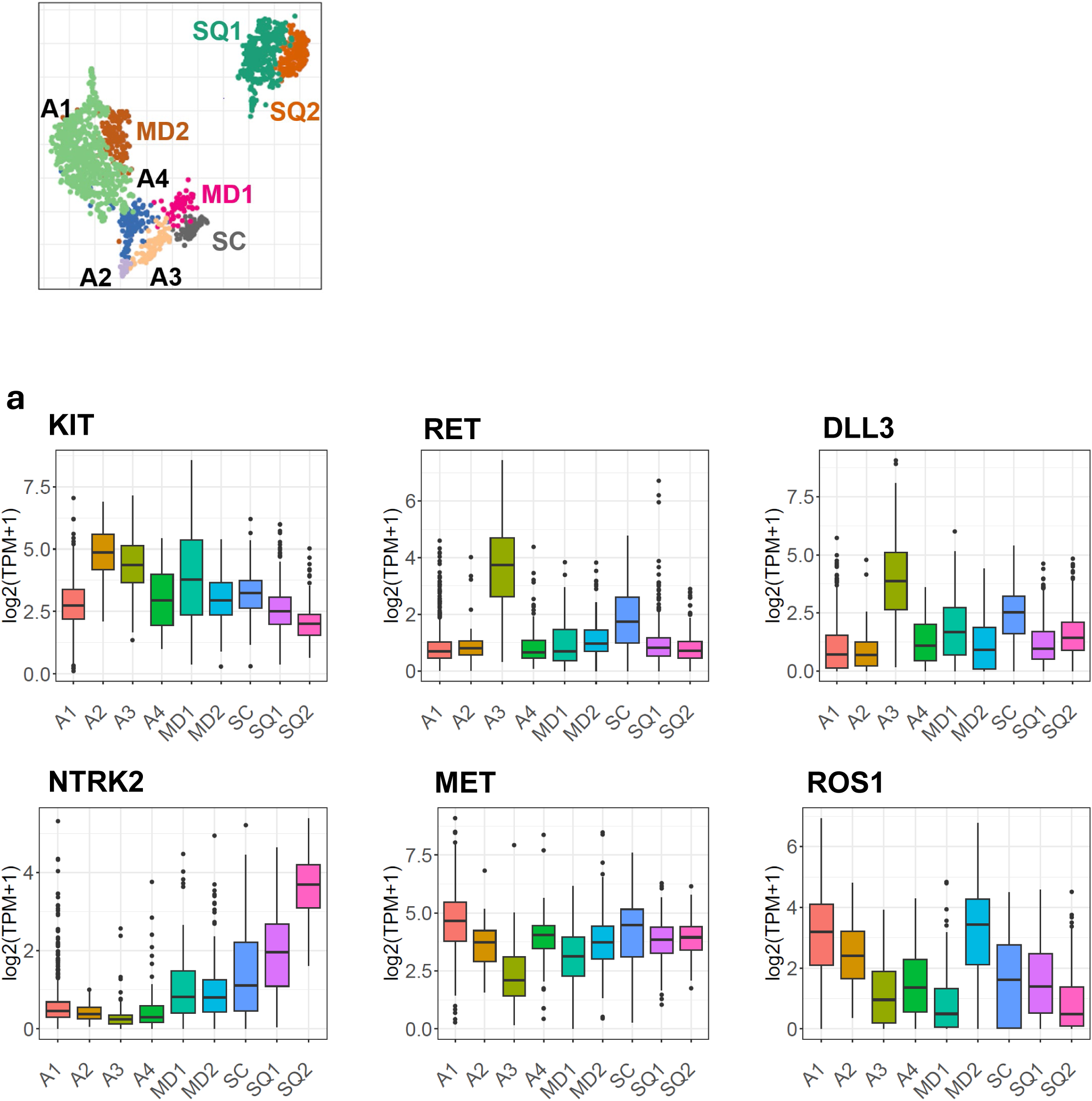
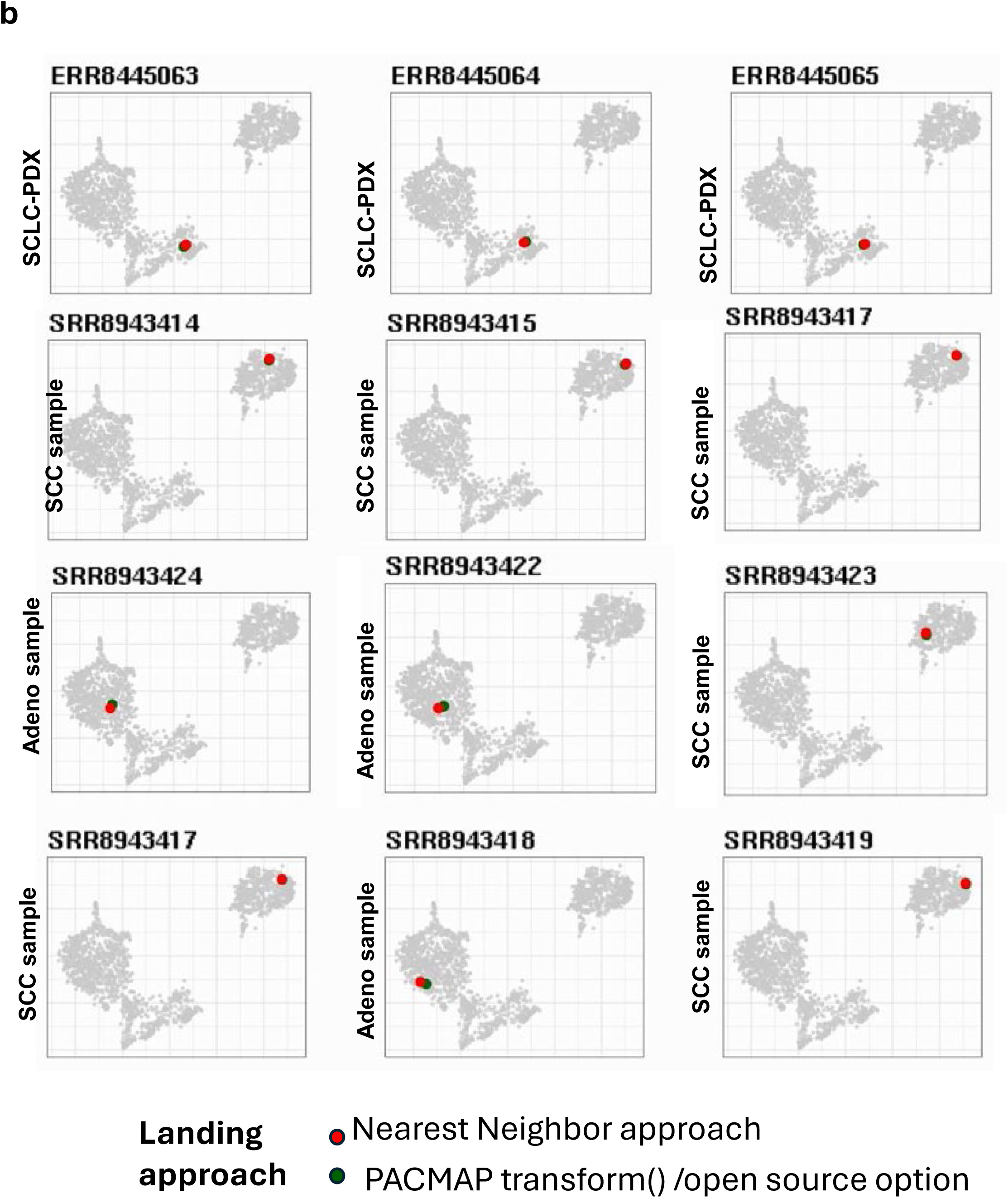
Cluster-specific expression of selected clinically relevant targets. (A) Boxplots showing expression of KIT, RET, DLL3, NTRK2, MET, and ROS1 across transcriptional clusters, highlighting cluster-specific enrichment of potential therapeutic targets. These distributions correspond to the spatial expression patterns shown in the main PaCMAP overlays in Figure 7A. Expression enrichment is interpreted as hypothesis-generating and does not establish functional dependency or therapeutic sensitivity. (B) Side-by-side comparison of two approaches for projecting patient samples onto the reference landscape. The red point indicates the nearest-neighbor landing approach, whereas the green point indicates the PaCMAP transform-based projection.

## Notes

### Competing Interest Statement

The authors have declared no competing interest.

### Summary of Updates

The manuscript has been revised. The revised manuscript now includes one new main figure comparing landscape clusters with established LUAD, LUSC, and SCLC subtype frameworks (Figure 3), a new main workflow schematic for projecting patient tumors or patient-derived xenografts onto the reference landscape (Figure 7C), and nine new supplementary panels, including dimensionality-reduction comparison and subsampling analyses (Figure S1B.C) and clinical, molecular, oncogenotype, mutation burden, tumor microenvironment, and neuroendocrine marker annotations across clusters (Figure S2E-K). We also added several new supplementary tables: pairwise survival comparisons across all nine clusters (Table S2), cluster-level clinical/molecular/tumor microenvironment association statistics (Table S5), concordance with established LUAD/LUSC/SCLC subtype frameworks (Table S6), and a candidate functional-testing roadmap for cluster-associated therapeutic hypotheses (Table S12). Together, these revisions sharpen the distinction between what the current atlas directly demonstrates and what it nominates for future validation.

